# Telomere-to-Telomere Genome Assembly Uncovers *Wolbachia*-Driven Sex-Specific Demography and Challenges Fisher’s Principle in a Sawfly

**DOI:** 10.1101/2024.12.12.628268

**Authors:** Mingpeng Zhang, Ruoyu Zhai, Gengyun Niu, Jiaqi Chen, Beibei Tan, Duo Wu, Guanliang Meng, Meicai Wei

## Abstract

*Wolbachia*, a widespread endosymbiotic bacterium, can reshape the evolutionary fates of its insect hosts by distorting reproduction and altering population dynamics. Despite extensive laboratory research, its long-term effects on host evolution in nature remain poorly understood, particularly regarding genetic mechanisms underlying changes in sex determination and reproduction. Here, we report the first telomere-to-telomere (T2T) genome assembly of the sawfly *Analcellicampa danfengensis* and the complete genome of it symbiotic *Wolbachia*. Comparative population genomics of six closely related *Analcellicampa* species revealed that *Wolbachia*-infected populations experience marked changes in sex-specific demography. While uninfected species maintain balanced genetic features between males and females, infected species show a persistent reduction in male effective population size alongside a stable or even growing female population, ultimately driving males toward extinction. Genomic scans identified positively selected genes associated with reproductive functions, sensory perception, neural development, and longevity, suggesting that *Wolbachia* manipulates critical host biological pathways to promote its transmission. These findings provide direct genomic evidence that *Wolbachia* acts as a powerful evolutionary force, reshaping host genomes in a way that disrupts Fisher’s principle, ultimately driving female-biased demography and the extinction of males at evolutionary timescales. This work provides deeper insights into host– endosymbiont coevolution and has important implications for evolutionary theory and pest management strategies.

## Introduction

Symbiosis in nature showcases the intricate and essential relationships between species. These interactions not only play a pivotal role in biological evolution but also significantly impact modern agriculture and ecosystem diversity^1-4^. Among the most extensively studied and evolutionarily remarkable examples of symbiosis is that of *Wolbachia pipientis*, a bacterium belonging to the order Rickettsiales within the α-proteobacteria^5^. This obligate intracellular symbiont infects millions of insects worldwide, with estimates suggesting that up to 66% of insect species harboring the infection^6^. Consequently, *Wolbachia* infections are considered one of the most widespread pandemics in the history of life, from a biodiversity perspective^7^. By exerting complex effects on the reproductive systems and population dynamics of its hosts, *Wolbachia* has become a focal point in contemporary biological research.

*Wolbachia* is renowned for its profound effects on the reproductive systems of insects. By inducing mechanisms such as cytoplasmic incompatibility, male killing, feminization, and parthenogenesis, *Wolbachia* significantly enhances its transmission and persistence within host populations^8^. These characteristics have made *Wolbachia* a promising candidate for pest control^9^ and for reducing the impact of arthropod-borne diseases^10^. While *Wolbachia* is primarily transmitted maternally through vertical transmission, it can also spread horizontally between species via predation, shared habitats, or symbiotic interactions^11^. These horizontal transfer between different host species, along with occasional infection loss, frequently cause discordance between *Wolbachia* and host phylogenies.

The microbiome frequently influences the range of host phenotypes across different taxa and environmental conditions, potentially playing a significant role in the evolution of hosts^12^. However, the full spectrum of the microbiome in shaping host phenotypic variance, evolutionary dynamics and demographic history is not yet fully understood. This gap in knowledge is also evident in *Wolbachia* research. Although the reproductive manipulation of *Wolbachia* on hosts was well-documented^1,8,13,14^, we lack a comprehensive genetic understanding of how it dynamically drives evolutionary changes at the population level or alters the evolutionary trajectory of its hosts during its symbiotic relationship in nature remain poorly understood, which impedes our understanding of *Wolbachia* outbreaks. Therefore, studying how *Wolbachia* affects hosts in epidemiological perspective, particularly in natural populations, becomes crucial. However, examples of host-*Wolbachia* interactions across different symbiotic stages within same or closely related species, especially in the wild, remain scarce.

In this study, we report the parasitism of *Wolbachia* in the sawfly *Analcellicampa* spp. (Hymenoptera, Tenthredinidae), whose larvae are fruit borer of *Cerasus* spp. (Rosaceae)^15^. This provides a unique opportunity to explore in detail how *Wolbachia* parasitism influences the evolution of this species, impacts host population demography and alters the host genome. We collected 89 individuals from six species across China and established a pipeline based on next-generation sequencing data to identify *Wolbachia* infections, discovering that three of these species were infected. We successfully assembled the first telomere-to-telomere (T2T) genome for *Analcellicampa danfengensis* (AD) along with the complete genome of its symbiotic *Wolbachia*. Utilizing these genomic resources, we investigated *Wolbachia* transmission characteristics in sawflies, applying an epidemiological perspective alongside genomic and population genetic approaches to assess its evolutionary impact. While previous research has predominantly focused on the origin, classification and reproductive manipulation of *Wolbachia*^13,16,17^, few studies have addressed its demographic effects and population evolutionary dynamics from a population genetics or epidemiological perspective. By examining the symbiotic relationship between *Wolbachia* and sawfly, we aim to provide new insights into the complexities of symbiosis, offering novel perspectives for the fields of ecology, evolutionary biology, and biological control. Furthermore, our findings may contribute to refining strategies for utilizing *Wolbachia* in the control of vector insect reproduction and transmission, ultimately enhancing pest management efforts.

## Results

### Assembly and Annotation of the *Analcellicampa danfengensis* Genome

Initially, we generated approximately 14.69 GB of Pacific Biosciences (PacBio) HiFi reads (∼68.89×), 54.80 GB of Oxford Nanopore (ONT) long reads (∼256.99×), 47.21 GB of Hi-C reads (∼221.39×), and 34.54 GB illumina paired-end reads (∼161.98×) (Table S1). Heterozygous k-mer pair coverage distributions from Smudgeplot^18^ revealed signals of diploidy (Figure S1A), consistent with previous studies^19^. The genome size of AD was estimated to be 213.24 Mb using GenomeScope^20^ (Figure S1B), close to the previously reported genome size of another sawfly species, *Orussidae abietinus*, at 201 Mb^19^.

We utilized various cutting-edge assembly software tools to construct the genome assembly. By almost every metric, Flye^21^ produced the best draft genome assembly, showing the most contiguous (N50) and high per-base quality among all assemblies, as indicated by its superior Benchmarking Universal Single-Copy Orthologs (BUSCO)^22^ score (Table S2). In contrast, LJA^23^ produced the lowest-quality assembly across all metrics, while Hifiasm^24^ in HiFi + ONT mode generated an intermediate-quality assembly. Therefore, we selected the Flye assembly for further scaffolding with Hi-C data and gap closure.

We used Hi-C data for scaffolding and then extended both ends of the scaffolds using HiFi reads to further elongate and link associated scaffolds (see Materials and Methods). This approach ultimately scaffolded the genome into 10 pseudo-chromosomes (Figure 1A and B), during which we also manually closed 77 gaps using the sequence-extension method. The final genome assembly was 211.14 Mb, featuring a contig N50 size of 22.39 Mb and a GC content of 41.99% (Table S2). The Hi-C contact map showed stronger interaction intensity along the diagonal compared to non-diagonal regions, with no significant noise outside the diagonal, suggesting a high-quality chromosome assembly (Figure 1A). The largest scaffold reached 57.04 Mb (Chr1), while the smallest was 13.31 Mb (Chr6). Seven scaffolds achieved telomere-to-telomere (T2T) assembly, though two scaffolds each still contained one unresolved gap each (Chr4:10537477-10537560 and Chr9:7931267-7931340) (Figure 1A). Assembly completeness was evaluated using the insecta-specific and hymenoptera-specific BUSCO database, showing 99.85% and 97.91% completeness, respectively, indicating robust genome integrity (Figure 1C and Table S2). Quality-control mapping of Illumina reads to the final assembly yielded an exceptionally high mapping rate of 99.83%. HiFi reads achieved a mapping rate of 99.93%. ONT reads achieved 98.83%, and RNA-seq data showed a mapping rate of 91.76%. These results further indicate the high quality of our genome assembly. Comparative assessments with AD and *Orussidae abietinus*^19^ genomes showed significant enhancements in the AD assembly, with a contig N50 approximately 17 times that of *Orussidae abietinus* and almost all gaps from previous versions filled (Figure S2). In summary, the assembly resulted in seven gap-free chromosomes, establishing AD as a high-quality chromosomal-level reference genome.

**Figure 1.**
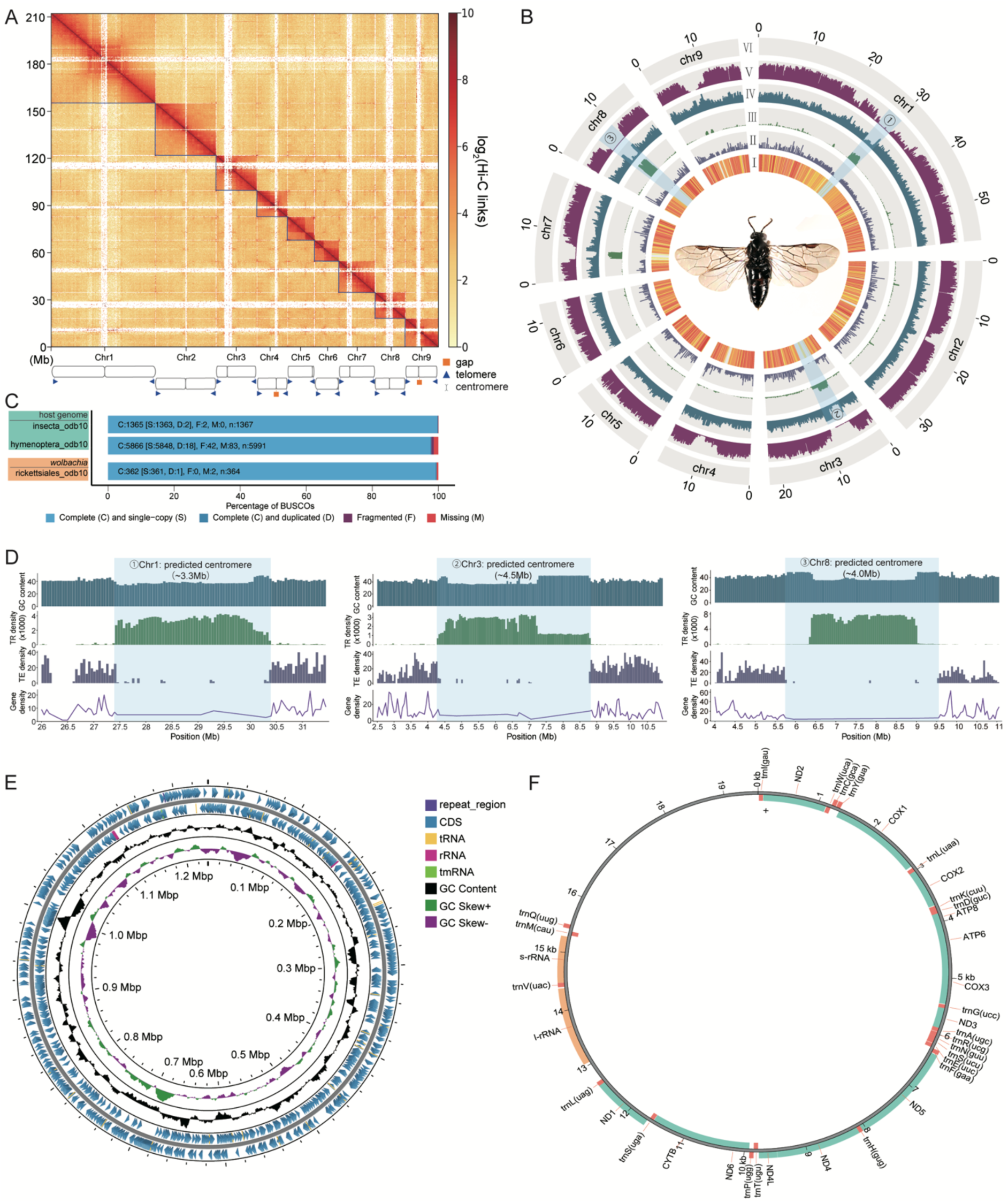
Assembly and key characterization of the *A.danfengensis* T2T genome, including centromeric characterization, symbiotic *Wolbachia* and mitochondrial DNA. A, Genome-wide Hi-C contact matrix of *A.danfengensis*. Red color intensity in the heatmap shows frequency of interaction between two loci at 25 kb resolution. B, Distribution of genomic features of the *A.danfengensis* genome. Tracks are aggregated in 100-kb bins as follows: I, gene density; II, TE density; III, TR density; IV, GC content; V, SNPs density; VI, the 9 chromosomes (chr1–chr9). C, BUSCO completeness scores for the host genome using insecta and hymenoptera databases, and for *Wolbachia* genome using rickettsiales database. D, features of the three largest centromeres: ① Chr1 (∼4 Mb); ② Chr3 (∼4 Mb); ③ Chr8 (∼3.5 Mb). E, Circular map of the *w*And genome. The outermost tracks 1 and 2 represent the positions of the CDS, tRNA, rRNA, and tmRNA genes on the positive and negative strands. Tracks 3 and 4 tracks show the GC content and GC skew, respectively. F, Features of the mitochondrial genome.

Moreover, we examined whether telomeres and centromeres were present in our assembled genome. The results showed that telomeric regions could be detected on both ends of nine chromosomes. The analysis predicted nine centromeric regions and identified 18 telomeres by recognizing the five-base telomeric repeat (CCTAA/TTAGG)^25^. Centromeric and telomeric regions are detailed in Tables S3 and S4, respectively. The approximate locations of the nine centromeric regions was estimated based on repeat density and the Hi-C interaction heatmap. We observed that most centromeres were positioned near to the middle parts of the chromosomes, with a few located at the chromosomes’ terminals, including Chr5 and Chr6 (Figure 1A, B and D, and Table S4). The centromeric regions ranged in size from 124.98 kb to 4.54 Mb, with an average length of 1.96 Mb (Table S4). The shortest centromeres are the telocentric, found on Chr5 and Chr6. Notably, “blank regions” observed in the Hi-C interaction map (Figure 1A) corresponded with a dense accumulation of tandem repeats, especially in the centromeric regions (Figure 1B and D), which is consistent with other T2T genome assemblies^26,27^. Centromeres vary greatly in their sequence organization among species^28^. Additionally, centromeric regions exhibit several notable common features. In AD, the centromeric regions are primarily composed of tandem repeats, accompanied by a reduction in GC content, a decrease in transposable element (TE) content, and a significantly lower gene density (Figure 1B). These distinct characteristics, compared to other genomic regions, further support the identification of centromeres.

A total of 9,569 protein-coding genes were predicted from the chromosomal-level genome of *Analcellicampa*. Of these, 4,716 genes were assigned Gene Ontology (GO) terms, 4,282 genes were assigned Kyoto Encyclopedia of Genes and Genomes (KEGG) pathways, and 6,787 genes contained Pfam domains. Additionally, 6,478 genes had BLASTN matches in the Swiss-Prot protein database. Repeat sequences constituted 32.42% of the AD genome, of which retrotransposons accounted for 12.29%, DNA transposons for 5.73%, and unclassified elements for 12.59%.

### Assembly and Annotation of the Endosymbiotic *Wolbachia* in sawfly

We generated a circular contig in our assembly (Figure S3), and sequence homology analysis confirmed it was *Wolbachia*. The circular structure indicates the completeness of the assembled sequence. The *Wolbachia* strain was detected in AD, thus we named it *w*And. The *w*And genome is 1,236,102 bp in size and has a GC content of 35.14% (Figure 1E and Table S5). Using Prokka software^29^, we annotated the genome, identifying a total of 1,166 genes were identified, 3 rRNA genes, 34 tRNA genes, and 1 tmRNA (Figure 1E). The BUSCO completeness scores of the *w*And genome indicated that it contained 361 complete and single-copy BUSCO groups, 1 complete and duplicated BUSCO groups, 0 fragmented BUSCO groups, and 2 missing BUSCO groups (Figure 1C). The BUSCO score of 99.5% for the *w*And genome was comparable to that of other circular, chromosome-level *Wolbachia* genomes in supergroup A^30^ (Table S6). The genomic features of *w*And, including coding sequences (CDS), tRNA, rRNA, and others, were visualized using the Proksee^31^ (Figure 1E). The genome exhibits the GC skew pattern typical of *Wolbachia* genomes^32,33^. The presence of an irregular GC skew in wAnd masks the identification of a specific origin of replication, indicating potential frequent genomic rearrangements^34^. In summary, we discovered *Wolbachia* parasitism in the sawfly and successfully assembled a high-quality *Wolbachia* genome.

We assembled a complete mitochondrial genome of 199,511 bp, encompassing all 13 protein-coding genes, 22 tRNA genes, 2 rRNA genes, and a control region (Figure 1F and Table S7). The control region is situated between *trnQ* and *trnI* and measures 4,064 bp in length.

### Resequencing Data Collection and High-Quality SNPs Construction

In this study, we collected 89 high-quality whole-genome resequencing data from across China (Figure 2A). Specifically, we obtained 30 AD, 26 *A.xanthosoma* (AX), 7 *A.acutiserrula* (AA), 13 *A.maculidorsatus* (AM), 4 *A.wui* (AW) and 9 *A.emei* (AE) for whole-genome sequencing and analysis (Table S8). We used two species *Monocellicampa pruni* and *Monocellicampa yangae* in closely related genus, as an outgroup. All 91 genomes were aligned against the AD reference genome. The lowest alignment rate was observed in the outgroup at 69.67%, while the average mapping rate for the *Analcellicampa* species was 89.07% (Table S8). Among the *Analcellicampa* species, the lowest alignment rate was observed in AE at 78.87%, while the highest alignment rates were observed in AD at 99.90%, followed by AX, with rates of 99.90% and 93.65%, respectively (Table S8). A total of 20,292,172 putative bi-allelic single-nucleotide polymorphisms (SNPs) were yielded and passed the filtering criteria across the 91 genomes for subsequent analyses, of which 11.83 million (39.62%) were intergenic, 6.83 million (22.86%) were intronic and 1.53 million (5.13%) were exonic (Table S9). We observed the highest number of SNPs per individual in the AE, with 3.39 million SNPs per individual, corresponding to ∼24.51% more SNPs than the average of 2.73 million (Table S8). To further assess the quality of the genetic variants, we calculated the transition-to-transversion (Ts/Tv) ratio, an indicator of potential sequencing error^35^. In our study, the Ts/Tv ratio for global populations SNPs was found to be 1.99, close to 2, indicating that the genetic variants identified are of high quality and their distribution is relatively balanced.

**Figure 2.**
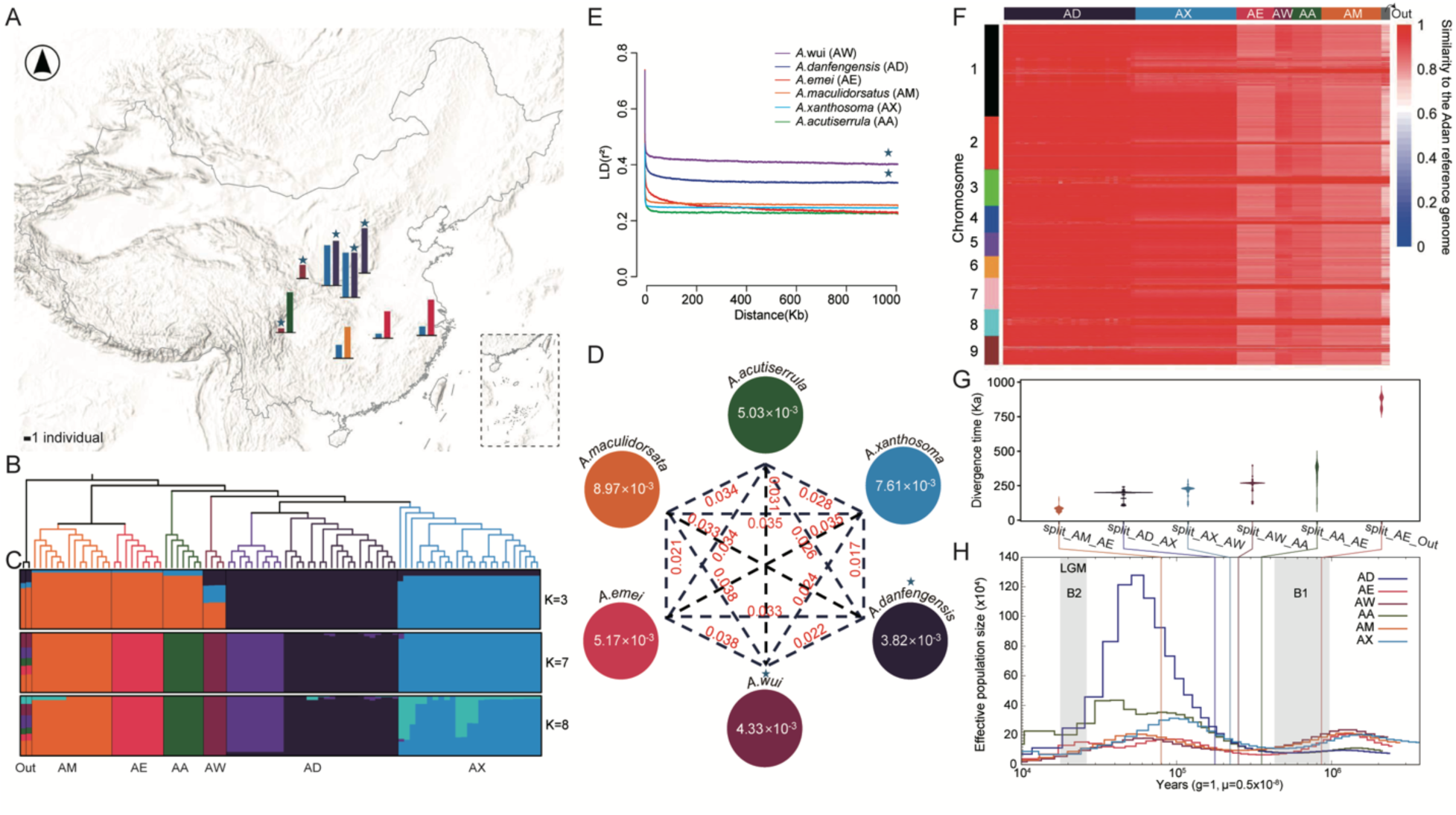
Population structure, phylogeny, and demographic history of *Analcellicampa*. A, Sampling sites. The bar charts indicate the species and number of samples collected at each sampling location. Colors correspond to those in panel D. B, Maximum-likelihood tree depicting the evolutionary relationships among genus *Analcellicampa* and outgroup. c ADMIXTURE analysis with K = 3, 7 and 8. Colors in each column represent ancestry proportion. D, Nucleotide diversity (π), nucleotide differences (dxy) across the six species. The value in each circle represents a measure of nucleotide diversity for each species; values in red on each line indicate pairwise population nucleotide differences between species. E, Patterns of LD (linkage disequilibrium) decay across the genome in different geographic populations. *r*^2^, Pearson’s correlation coefficient. F, Genomic similarity of six species of *Analcellicampa* to the AD reference genome. Chromosomes are indicated by different colors along the left y axis. Identical score (IS) values are shown for SNPs within each 50-kb window across the genome. G, Estimated split times between each species after 50 bootstraps. The widths indicate probability densities. H, Dynamic of *N*_e_ inferred by PSMC. The vertical lines indicate divergence times corresponding to those shown in G. The marked pentagrams indicate species fully infected with *Wolbachia*.

### Population characterization and evolutionary history of sawfly

To decipher genetic relationships among *Analcellicampa* sawfly populations, we constructed a Maximum Likelihood (ML) tree and performed principal component analysis (PCA) on the above 91 individuals using autosomal SNPs. Our phylogenetic and genetic clustering analyses resolved six genetic groups (Figure 2B and Figure S4), which corresponded to species classifications based on morphological characteristics (Figure S5 and S6). Using *Monocellicampa* species as outgroup, the phylogenetic analysis revealed that individuals from the same population clustered together, with AE and AM forming a clade distinct from the other genetic lineages in the ML tree (Figure 2B). Higher π values observed in AE and AM (with the exception of AX) reflected their rich genetic diversity (Figure 2D), potentially indicating that species from South and East China have more ancient origins compared to other populations (Figure 2A). Species from the northern and western regions clustered together, with each species forming stable genetic clusters (Figure 2A and B). Population structure inferred by ADMIXTURE^36^ also aligned with the six genetic groups when the optimal K = 7, with AD further subdivided into two subgroups corresponding to their geographical distribution (Figure 2C and S7). Additionally, when K values ranged from 2 to 6, the ancestry component of AE and AM accounted for the largest proportion in the outgroup (Figure S7), supporting the previous finding suggested by π values that populations from South and East China have more ancient origins compared to other populations.

Notably, AX and AD formed a sister clade (Figure 2B), with their dxy value being the lowest among all populations (0.017, Figure 2D). The geographic ranges of the two species overlapped, though AX had a broader distribution, with a few individuals still found in Hunan and eastern regions (Figure 2A and Table S7). AX exhibited the second-highest nucleotide diversity (π = 7.61 × 10^-3^), following AM (π = 8.97 × 10^-3^), while AD and AW showed the lowest nucleotide diversity (Figure 2D). Linkage disequilibrium (LD) decay analysis showed slower decline in LD for AW and AD (Figure 2E), suggesting smaller population sizes and further supporting their lower nucleotide diversity. Identity Score (IS) analysis demonstrated a high level of genomic similarity between AX and AD (Figure 2F). The dxy between AD and other species was approximately 0.03, with genomic IS reaching at least 0.6, indicating a close genetic similarity among populations and providing a solid foundation for our subsequent population genetic analyses using AD as the reference genome.

We also examined possible gene flow between species using TreeMix (Figure S8 and S9), which calculates a phylogeny of populations based on shared drift and tests whether migration edges (i.e., introgression) improve the model fit. The overall topology of the tree was consistent with that of the ML tree (Figure 2B), confirming the reliability of this tree’s topological structure. Likelihood improvements declined after four migration edges (Figure S8), so we presented results using this value (Figure S9), with 100 bootstraps performed. At four migration edges, TreeMix identified evidence for low levels of gene flow (migration weight < 0.05) from AA into AX, AE into AD, and a higher level of gene flow (∼0.2) from AD into outgroup (Figure S9).

Furthermore, we explored the demographic history of the sawflies by using site-frequency spectrum (SFS) via momi2^37^ and inferred changes in effective population size (*Ne*) over time for each population with the pairwise sequential Markovian coalescent (PSMC) model^38^. Since low levels of gene flow have negligible impact on timing inference^39,40^ and our study primarily focused on changes in *Ne* across species, we tested two models—one assuming constant population size and another allowing for variable population size—to seek the best one with lowest AIC value (see Materials and Methods and Figure S10). Based on this best-fit model (Figure S10), we inferred that the most recent common ancestor of the genus *Analcellicampa* diverged approximately 857,445 years ago (95% CI, 844.17 to 870.72 Kya). The basal clade within sawflies, containing AE and AM, separated from other lineages 368,609 years ago (95% CI, 313.15 to 384.07 Kya). Within *Analcellicampa*, the most recent divergence occurred between AE and AM at 79,678 years ago (95% CI, 74.39 to 84.97 Kya), followed by AD and AX at 185,866 years ago (95% CI, 174.66 to 197.07 Kya) (Figure 2G and Table S10). In the optimal model, we allowed population size to vary from the Last Glacial Maximum to the present. All populations showed a decline in *Ne*, though AW exhibited the fastest decline, followed by AD. This result aligns with the previous LD decay analysis, where these two populations demonstrated the slowest LD decay (Figure 2E).

The demographic history of the six species in our study was first inferred by analyzing the whole-genome sequence using Pairwise Sequentially Markovian Coalescent (PSMC) model^38^ (Figure 2H). The inferred demographic histories of these six species spanned from approximately 10 million years ago (Mya) to 10,000 years ago (Kya). Since all species analyzed here are younger than 10 million years, the inferred demographic dynamics likely reflect an ancestral form with a potentially different geographic distribution (Figure 2H). All sawfly populations experienced two population bottlenecks, one around 0.4 Mya (Bottleneck 1, B1) and another near 20 Kya (B2) (Figure 2H). Correlating species divergence times with *Ne* dynamics, we observed that the formation of the *Analcellicampa* genus occurred approximately 857 Kya, coinciding with the early phase of B1 (Figure 2G and H). The sawfly *Analcellicampa*, being monophagous with their larvae that exclusively parasitize cherry fruits (*Cerasus* spp.), likely experienced a founder effect at this bottleneck stage. Following B1, *Analcellicampa* underwent rapid radiation, accompanied by an increase in *Ne*. Over a span of approximately 300,000 years, six distinct species emerged, among which AD experienced a notable Ne expansion post-divergence. During the Last Glacial Maximum (LGM, ∼ 20 Kya), all populations experienced a decline in *Ne*, leading to the second bottleneck (B2).

To sum up, our results provided a comprehensive overview of the evolutionary history of *Analcellicampa*. The earliest emergence of the genus was around 857 Kya, originating in the southwestern region and subsequently spreading northeastward, resulting in six distinct *Analcellicampa* species in China. The most widely distributed species was AX, which diverged from its sibling species AD approximately 185 Kya. We did not observe any introgression events from AD, the species for which we constructed the reference genome, into other sawfly species. Overall, despite ∼857 thousand years of divergence, these populations maintain relatively high similarity, with IS values above 0.6.

### Discordance between phylogenetic relationships of the sawfly and its symbiotic *Wolbachia*

The transmission modes of *Wolbachia* play a crucial role in its spread and coevolution with hosts^41,42^. Our comprehensive genomic dataset allowed us to explore *Wolbachia* evolutionary dynamics within host populations, particularly to assess the extent of intraspecific horizontal transmission. Through our *Wolbachia* detection pipeline (See “Materials and Methods”), we observed infections in four sawfly species: AW, AM, AX and AD (Table S8). AW and AD were fully infected, while AM and AX showed partial infection, with AM having an infection rate of 3/13 among sampled individuals and AX exhibiting an infection rate of 1/26. Maximum likelihood phylogenetic trees of these *Wolbachia* strains classified them into three genetic groups: the first group parasitized only AW; the second included two individuals from AM; and the third encompassed all AD, one individual from AM, and the single *Wolbachia* strain found in AX (Figure 3). The *Wolbachia* strains found in AW form a distinct early-diverging lineage in the phylogenetic tree, suggesting a long independent evolutionary history of this host-symbiont partnership (Figure 3). The *Wolbachia* strains from AM were resolved into two distinct clades, while those from AD formed several closely related terminal clades with a lower genetic diversity (Type 1: 4.95 × 10^-3^ *vs*. Type 3: 2.05 × 10^-4^) and relatively fast LD decay (Figure S11). Two *Wolbachia* strains from AM formed a sister group relationship with the AD-associated strains, suggesting a shared evolutionary history between these infections. The single *Wolbachia* strain detected in AX was nested within the AD-associated clade, with both species sharing overlapping habitats. This relationship suggests a potential recent horizontal transfer from AD to AX, likely facilitated by ecological factors such as shared habitats and host density, aligning with the broader understanding that environmental context plays a critical role in host-symbiont evolution^43^.

**Figure 3.**
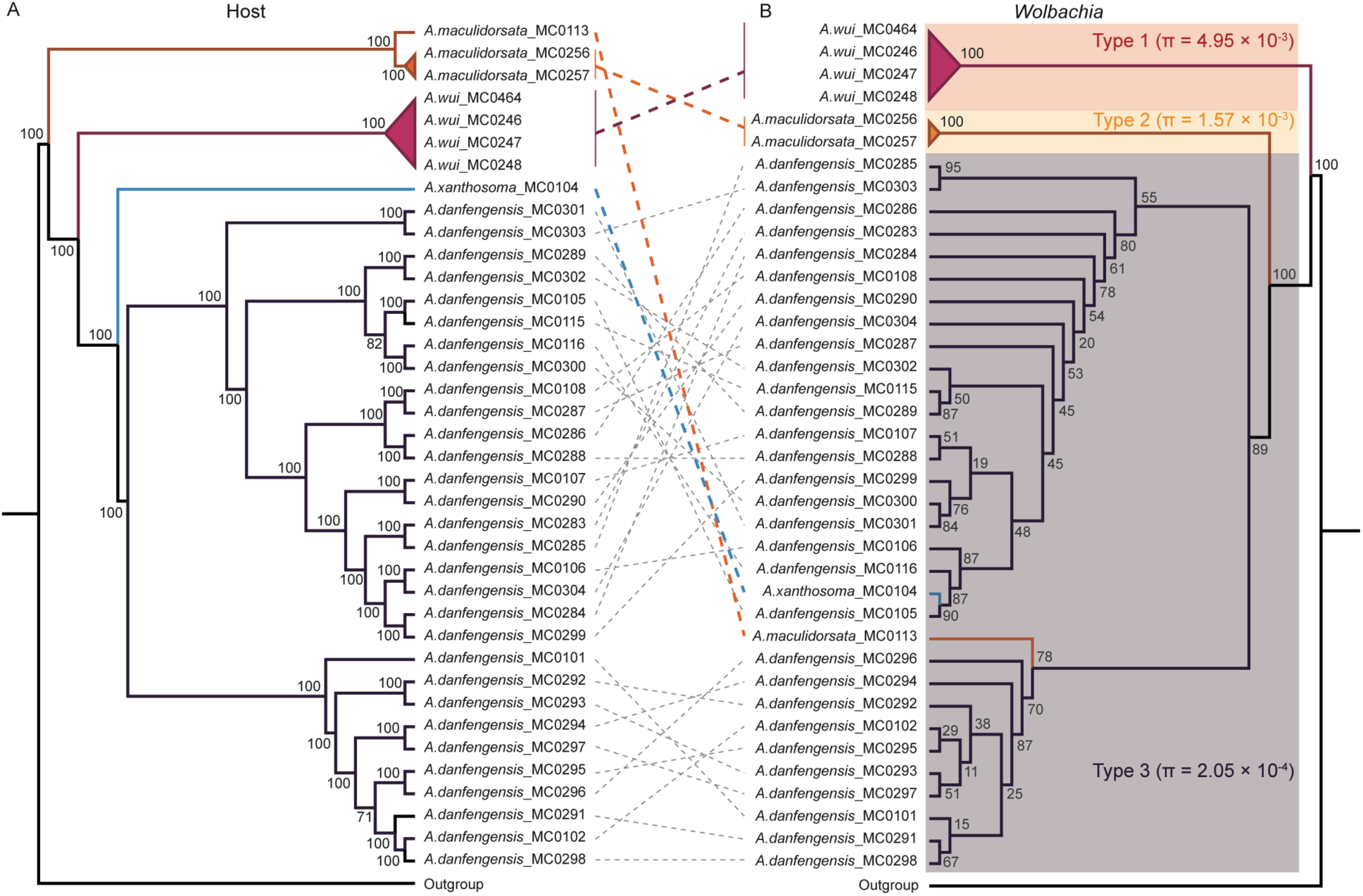
Phylogenetic relationships of *Wolbachia*-infected host and their *Wolbachia*. A, Phylogeny of host. B, Phylogeny of *Wolbachia*. The numbers at the major branches are bootstrap values.

When comparing host phylogeny with that of symbiotic *Wolbachia*, we found that the overall topologies of the host and *Wolbachia* trees were not fully congruent. However, some clusters maintained congruent phylogenetic relationships. For example, the phylogenetic patterns between AW and its associated *Wolbachia* showed strong congruence, suggesting stable host-symbiont associations likely due to long-term coevolution. The distribution of *Wolbachia* in AD showed partial congruence with host phylogeny, with both host and symbiont phylogenies revealing two main clusters that correspond to geographical distribution. However, within these clusters, the detailed phylogenetic relationships between hosts and their *Wolbachia* were discordant; For instance, while *A.danfengensis*_MC0301 and *A.danfengensis*_MC0303 formed sister clades in the host phylogeny (Figure 3), their associated *Wolbachia* strains showed greater phylogenetic divergence–this pattern was commonly observed within AD. These phylogenetic patterns are consistent with multiple modes of *Wolbachia* acquisition described in previous studies^11^, including vertical inheritance during host speciation, hybridization and gene flow between closely related hosts, and horizontal transmission events. In our study, these phenomena also reflect the existence of significant hybridization and horizontal transfer events.

### *Wolbachia* Associates with Sex-specific Demographic Changes in Sawfly Evolution

PSMC^38^ was used to infer historical changes in effective population size (*Ne*). The most dramatic demographic changes occurred between 20–200 Kya (Figure 2H), although the timing of population expansions and contractions varied among species. Interestingly, we observed significant differences between male and female AD during this period (Figure 4). To confirm this result, we inferred the demographic history of AD from three geographically distinct populations and performed 100 bootstrap replicates. The results remained robust, consistently showing significant differences in *Ne* between males and females during the 20–200 Kya period. Specifically, female *Ne* increased sharply around 200 Kya, peaking around 70–80 Kya, where female *Ne* was 13 times larger than that of males (13 × 10^5^ *vs.* 1 × 10^5^), before declining during the LGM to approximately 1 × 10^5^ (Figure 4A). In contrast, the male *Ne* remained relatively stable throughout this period, with a slight decline during the LGM.

**Figure 4.**
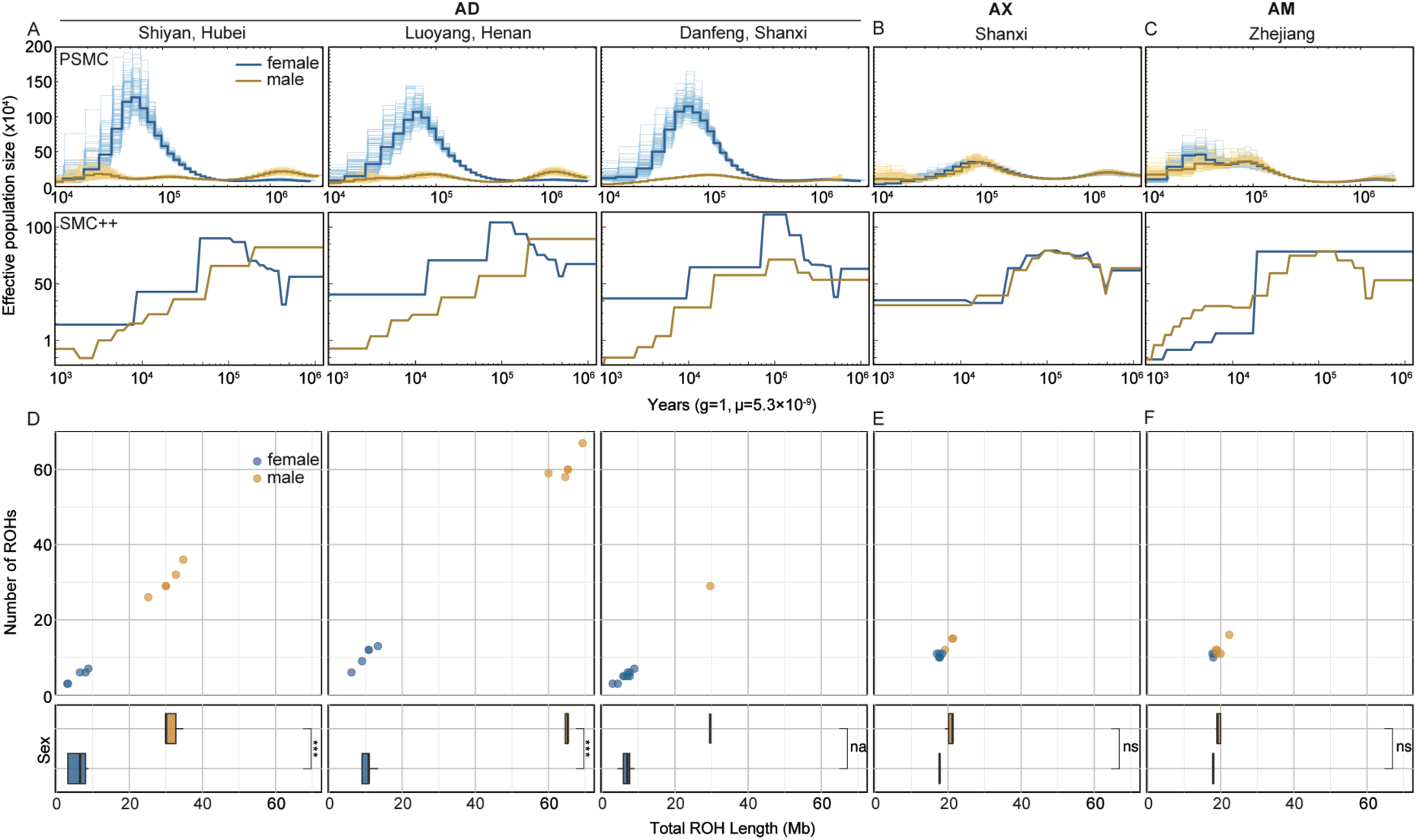
Demographic changes over time for female and male in populations fully infected with *Wolbachia* and those with limited *Wolbachia* infection, estimated using PSMC and SMC++. A, *N_e_* dynamics of females and males in three geographically distinct AD. B and C, *N_e_* dynamic of females and males in AX and AM, with *Wolbachia*-uninfected individuals used for the analysis. D, E, and F, Relationship between the number of ROH segments and the total ROH length in populations corresponding to those analyzed in panels A, B, and C. *** indicates *p* < 0.001; ns indicates *p* > 0.05; na indicates insufficient sample size, making statistical testing unavailable.

PSMC analyses based on re-sequenced data (Figure 4) enabled us to compare the demographic histories across the different populations. We included AX and AM as comparison groups (Figure 4B and C), as these populations contained predominantly uninfected individuals with only a small proportion showing recent *Wolbachia* infection. Using data from uninfected individuals, our analyses revealed highly similar demographic histories between males and females within these populations. Notably, the *Ne* of these uninfected individuals prior to 10,000 years ago were comparable to those observed for male AD (Figure 4A, B and C).

PSMC results become less reliable for recent timeframes (within the last 10,000 years) due to reduced inference power^38^. To address this limitation, we employed the SMC++^44^, which enable more precise inference of population history within the last 10,000 years. The demographic trajectories inferred by SMC++ prior to 10,000 years ago were largely consistent with the PSMC results, confirming an increased *Ne* of AD females around 70–80 Kya. However, SMC++ revealed that within the past 10,000 years, female populations remained stable, while male *Ne* experienced a dramatic decline, decreasing from approximately 10,000 to fewer than 1,000 individuals, indicating that the male population was on the verge of extinction. In contrast, populations (AX and AM) exhibited consistent demographic trajectories between males and females throughout their entire evolutionary history, despite differences in the range of *Ne* changes of two species. The size range and number of runs of homozygosity (ROH) fragments >700 kb were analyzed for male and female samples (Figure 4D, E and F). In the AD population, the total ROH length in males was significantly greater than that in females (approximately 3-6 times, based on mean values; Figure 4D). This result is likely due to a lower effective population size and a more concentrated genetic contribution among AD males.

Notably, a significant sex-biased *Ne* pattern was also observed in another fully *Wolbachia*-infected population, AW, where we sampled only four individuals (three females and one male, Table S8). The three females displayed similar demographic trajectories. For the male, PSMC could only infer *Ne* changes within the last 10,000 years, with *Ne* peaking around 2,000 individuals followed by a recent sharp decline (Figure S12). The inability to infer male *Ne* trajectory beyond 10,000 years ago likely reflects the combined effects of small population size and genetic drift, which can obscure signals of earlier coalescent events^45^. Given AW’s long association with *Wolbachia*, its males likely maintained a low *Ne* for an extended period. This prolonged low *Ne*, exacerbated by genetic drift, likely limits PSMC’s ability to detect earlier population histories^46^. As only one male AW individual was sampled, SMC++ analysis could not be conducted. Although we cannot directly infer the early *Ne* of AW males using PSMC, the current results indirectly suggest that this male population has been at a very low *Ne* for a long time. Additionally, over a decade of sample collection revealed significant sex ratio imbalances in AD and AW populations. Considerable effort was made to collect male AD individuals, while AW males were extremely scarce in the wild, making the single collected male sample exceptionally valuable.

The momi2 results further supported the population declines observed in AD and AW. We tested two models: one assuming constant *Ne* and another allowing for variable *Ne*. The variable *Ne* model, yielding the lowest AIC, demonstrated that *Wolbachia*-infected AD and AW experienced the fastest declines in *Ne*, and their current population sizes remain at low levels (Figure S10 and Table S10).

In summary, our analyses reveal complex temporal changes in host population dynamics associated with *Wolbachia* infection. While previous studies have largely focused on *Wolbachia*’s male-killing effects and their role in causing sex ratio imbalances, our findings in AD revealed a more complex process regarding sex distortion. Female *Ne* increased sharply around 70-200 Kya, followed by a decline during the LGM, whereas the decline of males was an ongoing process and a significant decline in male *Ne* occurred within the last 20,000 years (Figure 4). This sex-specific pattern indicates that the observed bias is not solely due to male reduction but involves an earlier expansion of female Ne and a prolonged male decline, reflecting complex host-symbiont interactions beyond simple male-killing. Additionally, by comparing populations with varying durations of *Wolbachia* infection, we found that infection length is closely related to the progression of male extinction. In AW, which had a relatively longer *Wolbachia* infection history, male *Ne* was at the lowest. In contrast, AD, with a relatively recent infection compared to AW, exhibited a less severe but ongoing reduction in *Ne*. This temporal gradient suggested that *Wolbachia*-infected populations experience a continual reduction in male numbers over time, making male disappearance appear inevitable. We did not observe a rapid increase in female Ne during the early stages of infection in AW as seen in AD. This difference may stem from the specific *Wolbachia* strain infecting AW, as different strains varied in their effects on host, such as distinct effects on immunity and reproductive traits^47-49^. Together, these results highlight how infection history and strain-specific factors drive distinct demographic trajectories in Wolbachia-infected populations.

### Genomic Signatures of *Wolbachia*-Induced Changes in AD

*Wolbachia* is known to manipulate host reproduction through male killing, feminization, parthenogenesis, and cytoplasmic incompatibility^8,50,51^, yet the genetic mechanisms underlying these effects remain poorly understood. Leveraging the unique system of sympatric sibling species—AD (fully infected) and AX (nearly uninfected)—with overlapping distributions and shared ancestry, we had a rare opportunity to explore how *Wolbachia* modifies host reproduction at the genetic level. By focusing on these populations, we minimized confounding variables, such as environmental effects, which are known to influence *Wolbachia*’s impact on sex determination^52,53^. This system, with its genomic similarity and robust sample sizes, allowed us to investigate *Wolbachia*-induced genetic modifications in an ecologically controlled context.

Specifically, we analyzed whole-genome sequences of AD (n=30) and AX (n=26) to investigate *Wolbachia*-induced modifications in the host genome. To identify the genomic region that presents signatures of natural selection associated with *Wolbachia*-coevolution, we performed whole-genome genetic differentiation analysis by estimating the cross-population composite likelihood ratio (XP-CLR) and differences in nucleotide diversity (π log_2_-ratio *Wolbachia*_positive/*Wolbachia*_negative) between AD and AX in 10-kb windows with a 5-kb sliding window along the genome. Applying a 5% threshold for maximum haplotype frequency difference (XP-CLR) and nucleotide diversity (log_2_-π ratio), we identified a total of 3.94 Mb genomic regions that covered all outliers (Figure 5A). These regions contained 287 annotated positively selected gene (PSG) candidates (Table S11).

**Figure 5.**
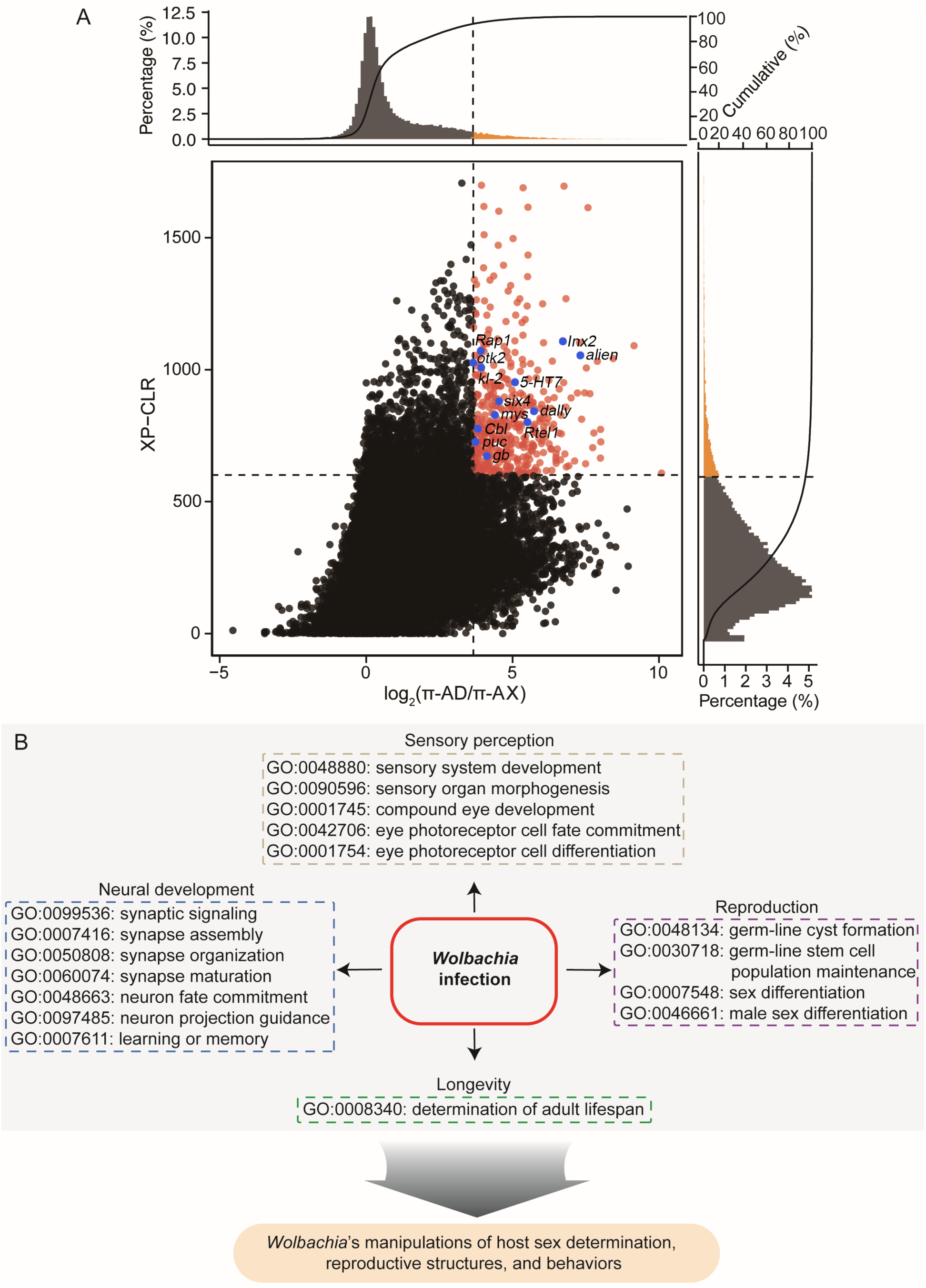
Selective sweep and enrichment analysis of the Wolbachia-infected AD. A, Distribution of the log_2_(π*-*_AD_/π*-*_AX_) and XP-CLR values were calculated in 10 kb sliding windows with 5 kb steps. The horizontal and vertical lines represent threshold lines of the top 5% of the XP-CLR and π ratio values, respectively. Points (red) located in the top right sector represent selective signatures in AD. Yellow and black bins in the histograms of XP-CLR (right) and π ratio (top) represent levels respectively higher and lower than the threshold line. B, Significant GO enrichment of PSGs associated with *Wolbachia*-induced reproductive manipulations.

Gene Ontology (GO) analysis of the PSGs revealed significant enrichment in categories associated with reproductive processes, sensory and neural development, among others (Table S12). Notably, the most highly enriched Gene Ontology (GO) terms were those associated with sensory system development (*P* = 3.0 × 10^-4^, Benjamini-Hochberg corrected) and compound eye development (*P* = 4.0 × 10^-4^) (Figure 5B). Enrichment was also observed in reproductive development categories, such as germ-line stem cell population maintenance (*P* = 1.3 × 10^-3^), germ-line cyst formation (*P* = 1.4 × 10^-2^), male sex differentiation (*P* = 1.4 × 10^-2^) and sex differentiation (*P* = 3.8 × 10^-2^) (Figure 5B). Additionally, neural development terms like synaptic signaling (2.7 × 10^-3^), synapse assembly (*P* = 9.9 × 10^-4^), synapse organization (*P*=1.3 × 10^-2^), regulation of synapse organization (*P* = 1.2 × 10^-2^), regulation of synapse structure or activity (*P* = 6.9 × 10^-3^), synapse organization (*P* = 1.3 × 10^-2^), neuron fate commitment (*P* = 2.7 × 10^-3^) and learning or memory (*P* = 1.2 × 10^-2^) were significantly enriched (Figure 5B). Notably, we also observed significant enrichment in the category of determination of adult lifespan (*P* = 4.4 × 10⁻^2^) (Figure 5B), suggesting potential selective pressures related to longevity and its influence on population dynamics under *Wolbachia* infection.

We identified several reproduction-related genes potentially linked to *Wolbachia*’s sex-manipulation mechanisms, corresponding to the significant demographic differences observed between males and females in our study. Among the reproductive GO terms we identified, the following genes were included: *mys*, *otk2*, *puc*, *Six4*, *alien*, *Cbl*, *Inx2*, *dally*, *Rap1*, and *Rtel1*. Among these, *Inx2* exhibited the strongest selection signal with XP-CLR=1297.74 and log_2_(π ratio)=4.85, while *puc* showed the lowest signal with XP-CLR=700.75 and log_2_(π ratio)=3.69 (Table S11).

Notably, *Six4*, which encodes a transcription factor essential for embryonic gonad formation, has been implicated in sex determination pathways. Its mammalian homolog regulates *Sry* expression, a gene essential for male development, and the loss of *Six4* function can result in the reversal of sex from male to female in mice^54^. This gene similarly contributes to *Wolbachia*-induced feminization of genetic males, as observed in some isopod (Crustacean) species and lepidopteran species^5^, where *Wolbachia* cause male embryos to develop as functional females. This process may underlie the increased female population numbers observed in our study.

Several of the above genes are associated with sperm formation, highlighting *Wolbachia*’s potential influence on male reproductive capabilities. For instance, *Rtel1*, a regulator of telomere elongation helicase 1, plays a role in male germline stem cells regulation^55^*. asun* is essential for spermatogenesis in *Drosophila*, with mutations leading to male sterility^64^. *Dcst2* is required for sperm-egg fusion, with knockout resulting in impaired fertilization in zebrafish^56^. Additionally, genes such as *Otk2*, linked to male genitalia development^57^, and *Inx2*, which encodes an innexin protein critical for gap junction communication during oogenesis, may influence oocyte development and fecundity^58^, underscoring their importance in the development of the male reproductive system.

Beyond the genes enriched in the aforementioned GO terms, we identified three additional reproduction-related genes, *kl-2* (XP-CLR = 978.52, log_2_(π ratio)=4.03), *5-HT7* (XP-CLR = 868.21, log_2_(π ratio)=4.88) and *gb* (XP-CLR = 810.20, log_2_(π ratio)=4.71). *kl-2* is a male fertility factor, is involved in flagella movements^59^. The remaining two genes are linked to mating behavior. *5-HT7*, a serotonin receptor, regulates sexual receptivity in virgin female *Drosophila melanogaster*, and its absence significantly reduces the mating rate of female flies^60^. Meanwhile, *gb (genderblind)*, encoding an amino acid transporter involved in glutamate secretion and synaptic transmission, was identified as a target of selection and is known to affect male courtship behavior in *Drosophila*^61^. Notably, a majority of these reproduction-related genes (8 out of 13) are located on Chr1 (Table S11), suggesting potential chromosomal clustering of selection signals.

These findings underscore the potential for *Wolbachia* infection to drive significant genomic changes in host genes related to reproduction, development, and behavior, highlighting the multifaceted impact of *Wolbachia* on host biology. By exerting selective pressures on critical biological processes, *Wolbachia* likely enhances its own transmission success^62^, while simultaneously altering host sex ratios and driving male extinction through complex genetic mechanisms.

## Discussion

Our findings reveal compelling evidence linking *Wolbachia* infection status to shifts in host sex ratio dynamics. Specifically, the geographic and temporal distribution of observed changes in host sex ratio dynamics—such as female-biased demography and reductions in male population size—corresponds closely to established mechanisms of *Wolbachia*-induced reproductive manipulation. Supporting evidence from controlled laboratory investigations has extensively documented *Wolbachia*’s capacity to reshape host reproductive biology through mechanisms such as male-killing, parthenogenesis induction, and behavioral alterations in mating systems^13,63^. Particularly compelling is our detection of strong selective signatures in genetic loci associated with reproduction, neural development, and sperm formation. These genomic regions correspond directly to phenotypic traits known to be targeted by *Wolbachia*’s manipulations, offering mechanistic support for *Wolbachia*-driven selection as a primary driver of the observed evolutionary shifts. The consistency between molecular signatures of selection and experimentally verified *Wolbachia*-host interactions^13,64^ strengthens the foundational understanding linking infection status to population-level reproductive dynamics. This integration of genomic, phenotypic, and experimental evidence provides robust support for the hypothesis that *Wolbachia* acts as a significant selective force, fundamentally shaping host evolutionary trajectories.

Moreover, alternative explanations, such as neutral demographic shifts or environmental factors, are less consistent with the observed patterns. Neutral demographic processes, such as genetic drift, population bottlenecks, or random migrations, typically result in stochastic genomic changes that are distributed randomly across the genome^65^. In contrast, we observe concentrated selection signals specifically enriched in functional categories tied to *Wolbachia*’s reproductive manipulations, such as genes associated with reproduction and mating behavior. This discrepancy suggests that the observed genomic patterns are more likely driven by directional selective pressures than by random demographic events. Similarly, environmental factors alone are insufficient to account for the congruence between *Wolbachia* infection rates and the persistent female-biased demography observed in AD and AX. Environmental variation typically produces diffuse and temporally variable effects, often fluctuating with changing ecological conditions^66,67^. By contrast, the observed patterns exhibit a more targeted and consistent influence, consistent with selective pressures rather than stochastic or transient factors. Collectively, these findings provide compelling, albeit indirect, evidence that *Wolbachia* acts as a major selective force, reshaping host reproductive dynamics over evolutionary timescales.

Surprisingly, our results reveal a remarkable deviation from the longstanding assumption that host populations stabilize at symmetrical sex ratios. Traditional theoretical frameworks, epitomized by Fisher’s principle^68,69^, posit that deviations from equal offspring investment generate countervailing selective pressures that restore equilibrium in sexually reproducing populations. However, the persistent, maternally inherited influence of *Wolbachia* challenges this paradigm, presenting an exception to the universality of Fisherian dynamics. Rather than converging on a stable balance, populations such as AD and AW, infected by *Wolbachia*, exhibit persistent, gender-biased evolutionary dynamics. In these populations, female-biased demography becomes progressively entrenched, while the evolutionary prospects of males steadily decline. This sustained imbalance disrupts the typical patterns of sex ratio equilibrium and the typical genetic contributions of both sexes, underscoring how microbial symbionts can fundamentally reshape host population structures and evolutionary trajectories over time.

Importantly, the mechanisms underlying these dynamics extend beyond well-documented effects such as cytoplasmic incompatibility or male-killing observed in laboratory settings or short-term natural infections^8,13,63^. By integrating T2T genome assembly, population genomics, and phylogeographic analyses, our study identifies signatures of positive selection on genes associated with reproduction, sensory perception, neural development, longevity, sperm formation, and other complex phenotypes. These genetic imprints offer a plausible mechanistic explanation for the persistent deviations from Fisher’s principle: when key genes influencing mate choice, gamete formation, and sexual behavior are reshaped by *Wolbachia*-driven selection, this equilibrium begins to erode. Over time, such modifications may recalibrate mating systems, alter sexual signaling, and increase the reproductive value of female progeny. While this does not preclude the involvement of other biological complexities, it provides a compelling framework for understanding how *Wolbachia* can undermine the self-correcting forces central to Fisherian dynamics. This disruption ultimately locks populations into sustained female-biased dynamics, fundamentally reshaping their evolutionary trajectories.

While other deviations from Fisher’s principle exist in nature, including phenomena such as local mate competition in structured populations (e.g., fig wasps where males compete within localized environments)^70^ or environment-dependent sex determination in reptiles like turtles and alligators, where temperature influences the offspring’s sex^71^. However, these instances are typically transient or context-specific, driven by short-term ecological pressures or localized conditions. In contrast, the Wolbachia-driven sex ratio imbalance we reveal appears fundamentally distinct due to its persistence and genetic basis. For example, *Wolbachia* infection in *Ostrinia* moths has led to male-killing^72^, and in certain Hymenoptera, parthenogenesis has eliminated male reproduction entirely^73^, showcasing the range of its reproductive manipulations. Rather than being a response to immediate environmental changes, this imbalance results from long-term, symbiont-mediated modifications to host genetics. Such sustained interactions alter the evolutionary trajectories of host populations in ways that are difficult to reverse. By reshaping key genetic pathways related to reproduction, behavior, and population dynamics^74^, *Wolbachia* demonstrates the ability to drive far-reaching and enduring shifts in sex ratios. This microbially driven imbalance challenges traditional models, which often focus on short-term ecological fluctuations, and underscores the profound evolutionary impact of microbial symbionts as architects of host genetic and demographic landscapes.

Beyond illuminating the fate of host reproduction, our results highlight *Wolbachia*’s remarkable ecological and evolutionary adaptability. Although vertical, maternal inheritance has long been considered the primary transmission route, the broad phylogenetic distribution of *Wolbachia* suggests that horizontal transfer also plays a pivotal role^43,75,76^. Our phylogenetic analyses reveal incongruence between host and *Wolbachia* lineages, indicating inter- and intra-specific transfers likely facilitated by ecological interactions such as shared habitats and predation^11,77^. For example, the close phylogenetic relationship between *Wolbachia* strains in AD and AX, which overlap geographically, suggests recent horizontal transfer. Such horizontal transmission strategies enable *Wolbachia* to colonize new host species, fostering novel coevolutionary relationships. Additionally, contrasting patterns of genetic diversity highlight different transmission dynamics: in long-established associations like AW, stable coevolution has produced high genetic diversity and slow LD decay, signaling a tightly integrated partnership, whereas in more recent infections, as seen in AD, AM, and AX, lower genetic diversity and frequent horizontal transfers mirror a dynamic phase of host-symbiont negotiation. These findings demonstrate *Wolbachia*’s diverse and adaptable transmission strategies, enabling its persistence across varied ecological contexts.

From a practical perspective, these findings carry substantial implications for the use of *Wolbachia*-based strategies in vector control and pest management. While the short-term goal of suppressing disease vectors or agricultural pests through mechanisms like cytoplasmic incompatibility is promising^78,79^, our data highlight the need for caution when considering the long-term evolutionary and ecological consequences. Persistent female bias in target populations could trap them in demographic cul-de-sacs, reducing overall genetic diversity and constraining their adaptive capacity in changing environments. Such genetic bottlenecks might initially seem beneficial for pest suppression but could lead to broader ecological disruptions, including niche vacuums that might be filled by more invasive or harmful species^80^. Furthermore, horizontal transmission of *Wolbachia* to non-target species—such as pollinators or other beneficial insects—poses additional risks, potentially destabilizing ecosystems. In an era of rapid environmental shifts and emerging pest challenges^81^, it is crucial to anticipate these long-term outcomes. A deeper understanding of *Wolbachia*’s genetic, demographic, and ecological impacts will help design more sustainable and responsible pest management strategies, minimizing risks while maximizing benefits. Recognizing the potential for unintended consequences in non-target populations is essential to ensure that these interventions achieve their goals without compromising broader ecosystem stability.

Looking ahead, addressing the interplay between host genetic selection, symbiont strain variability, and ecological pressures is crucial to advancing our understanding of *Wolbachia*’s evolutionary role. Approaches such as experimental evolution^82^ and long-term ecological monitoring^83^ would directly test the mechanisms proposed in this study, bridging genomic insights with broader evolutionary principles.

In sum, our findings depict *Wolbachia* as more than a reproductive parasite or mutualist. It emerges as a potent evolutionary force capable of reshaping sex ratios, sculpting genomic landscapes, and fundamentally altering host population histories. Far from being an evolutionary curiosity, this scenario underscores the pervasive influence of microbial symbionts, challenging traditional models and broadening the conceptual scope of evolutionary biology. By demonstrating that *Wolbachia* can drive persistent deviations from Fisher’s principle, establish female-biased population structures, and potentially set the stage for parthenogenesis, our findings highlight the profound and lasting role of microbial symbionts in reshaping host evolution. Rather than being a historical anomaly, these interactions reveal the enduring importance of co-adaptation and conflict in shaping biodiversity. Ultimately, this work emphasizes the necessity of integrating microbial symbionts into the broader narrative of life’s complexity and adaptive scope, redefining evolutionary trajectories over deep timescales.

## Materials and Methods

### Sample Collection and Sequencing

We collected samples from six species of the genus *Analcellicampa* (AD, AX, AA, AM, AE, AW). Two previously sequenced and published species from the sister genus *Monocellicampa*^84^ were used as outgroup references. For AD, we performed PacBio SMRTbell library construction and sequencing using the PacBio Sequel II platform, with data quality evaluated through CCS v6.0.0 software (https://github.com/pacificbiosciences/unanimity). High-accuracy circular consensus sequencing (CCS) reads were obtained for downstream analysis, yielding a total of 14.69 GB of HiFi data. Hi-C data was generated on MGI sequencing platform to assist in genome assembly. Additionally, 54.80 GB of ultra-long read data was obtained from Oxford Nanopore Technology (ONT) sequencing. The remaining samples were sequenced using Illumina paired-end sequencing, with achieving depths exceeding 20× (Table S8).

### Genome Survey and Assembly

Genomic sequences were characterized for size, heterozygosity and repetitiveness through k-mer frequency analysis (k = 21) using KMC3 [v3.2.1]^85^ and GenomeScope 2.0^18^. To ensure reads are reliable, Illumina paired-end sequenced raw reads for the genomic survey were first filtered using the Fastp v.0.20.0 preprocessor (set to default parameters)^86^. Heterozygous k-mer pairs were then analyzed using Smudgeplot^18^ to estimate ploidy levels and infer genomic complexity.

The Nanopore sequencing platform was used for the Ultra-long sequencing of DNA samples. The failed reads were removed from the raw data and We quality-filtered male and female ONT reads using NanoFilt [v2.5.0]^87^ software was used to filter the fragments <10 kb. The obtained pass reads were used for subsequent analyses. The joint sequence was filtered using Porechop v0.2.4 software (https://github.com/rrwick/Porechop), reads with retention lengths ≥30 kb and mean read quality scores >90% were used for assembly.

To maximize the quality of the final assembly with these data, we generated and compared several draft genome assemblies using four methods: Flye [v2.9.2]^21^, hifiasm [v0.18.2]^24^ and LJA [v0.2]^23^. Each assembly was assessed the contiguity and completeness using the program compleasm^88^ against the insecta ortholog database (Table S2). Based on these evaluations, we selected the highest quality genome assembly (the Flye assembly; Table S2) to proceed with Hi-C scaffolding. we mapped Hi-C reads to the draft assembly using BWA [v0.7.17]^89^ and scaffolds were constructed using YaHS [v1.2a.1]^90^, producing 232 scaffolds. HiFi reads were compressed following a process similar to Canu^91^ and used to extend scaffolds larger than 1 Mb with custom scripts. During each round of extension, we performed a BLAST search of the extended sequences against the assembly to check for overlaps with the assembled scaffolds. If an overlap was consistently and stably detected at the ends of two scaffolds, the two scaffolds were connected; otherwise, the extension continued until it could no longer proceed. Telomere sequences were also extended as much as possible using this method. Then, assembly underwent two rounds of additional polishing with NextPolish^92^ using resequencing data. Using this method, we ultimately yielded nine chromosome-scale scaffolds with no apparent large-scale misassemblies. The final Hi-C contact map was visualized with Juicebox [v1.11]^93^ to identify and correct potential misassemblies.

In addition, the complete *Wolbachia* genome (1,236,102 bp) was assembled using the Flye assembler and polished as part of the aforementioned process. The *Wolbachia* genome quality was evaluated using BUSCO scores with the rickettsiales_odb10 database. We annotated the *Wolbachia* genome with Prokka [v1.1.1]^29^. We assembled and annotated the mitochondrial genome (199,511 bp) and using the MitoHiFi [v3.2]^94^.

### Identification of telomeres and centromeres

We employed quarTeT^95^ to predict both telomeres and centromeres. Telomeres were predicted using quarTeT TeloExplorer module with the parameters: ‘-c other -m 100’. Centromeres were predicted using quarTeT CentroMiner module with the parameters: *‘-n 100 -m 200 -s 0.8 -d 10 -e 0.00001 -g 50000 -i 100000 --trf 2 7 7 80 10 50 -r 3 --TE EDTA.TEanno.gff3*’. Additionally, centromFind, with default parameters, was used to further validate centromere positions. These tools utilize complementary approaches for centromere prediction. CentroMiner identifies centromere candidates based on genome data and transposable element (TE) annotation, focusing on tandem repeats and surrounding TE regions. In contrast, CentromFind uses Hi-C matrices to detect centromeres by identifying regions with low interaction frequencies, which are indicative of centromeric repetitive sequences. The integrated results from the quarTeT CentroMiner module and centromFind. When both tools identified centromeres, their intersection was taken. If only one tool produced predictions, those results were directly used.

### Annotation of Repetitive Sequences

De novo repeat library prediction and homology-based comparisons were used for the annotation of repetitive sequences. We used RepeatModeler [v2.0.2] (https://github.com/Dfam-consortium/RepeatModeler) with default parameters to build the de novo repeat library. LTR_FINDER [v1.70]^96^ and LTR_retriever [v2.9.0]^97^, both with default parameters, were applied to identify long terminal repeat (LTR) sequences in the genome. Annotation and masking were performed using RepeatMasker [v4.1.5]^98^, following the pipeline (https://darencard.net/blog/2022-07-09-genome-repeat-annotation/). Specifically, the genome was annotated and masked in a stepwise manner using RepeatMasker: simple repeats, were annotated first, followed by curated repeat elements from existing databases. Next, known species-specific repeats identified from the sixth iteration of RepeatModeler were annotated, and finally, unknown species-specific repeats from the same cycle were incorporated.

### Genome Annotation

The genomic region containing repetitive sequences was masked and utilized for subsequent analyses. We employed three complementary strategies to predict protein-coding genes: homology-based prediction, transcriptome-based prediction, and *ab initio* gene prediction. For the transcriptome-based approach, we first mapped clean transcriptome sequences to the assembled genome using HISAT2 [v2.1.0]^99^. Transcript structures were then reconstructed with StringTie [v2.2.1]^100^, and coding regions were identified using TransDecoder [v5.5.0] (https://github.com/TransDecoder/TransDecoder). To refine these predictions, we used BLAST and HMMER to compare TransDecoder’s predicted coding sequences with known protein databases. The results from these homology searches were integrated into the transcriptome-based gene predictions to improve accuracy. For homology-based annotation, we used Miniprot [v0.13]^101^ to align homologous protein sequences (Arthropoda) to the genome, enabling the inference of unknown gene structures. This approach allowed us to predict gene functions, domains, and other characteristics by comparing the target genome with known homologous protein sequences. For *ab initio* gene prediction, we utilized Braker3 [v3.0.6]^102^ and Augustus [v3.5.0]^103^ with default parameters to perform gene prediction on the repeat-masked genome assembly. During the Braker3 run, we incorporated protein homology information from OrthoDB of arthropoda to enhance prediction accuracy. Finally, we integrated gene models from the different prediction methods using EvidenceModeler [v2.1.0]^104^ to produce a consolidated set of gene predictions. Following gene prediction, we performed functional annotation by matching predicted genes against various databases. Specifically, we used BLASTP [v2.12.0] with a threshold of 1e-5 to compare predicted genes against the SwissProt protein database. Additionally, we employed the eggNOG-mapper web server (http://eggnog-mapper.embl.de/) for functional annotation, including Gene Ontology (GO) terms, KEGG pathway annotations, and Pfam domain annotations.

### Preliminary analysis of resequencing data to identify *Wolbachia* Infection using HAYSTAC

*Wolbachia*, like most endosymbionts, cannot be cultured independently^16^. Consequently, even when enrichment protocols are employed, host DNA is typically co-sequenced with *Wolbachia* in the same experiment. This presents challenges related to sequencing coverage and potential contamination, as the host genome is often two to three orders of magnitude larger than the *Wolbachia* genome. Therefore, implementing a bioinformatics pipeline is crucial for detecting *Wolbachia* within host resequencing datasets. Here, we screened 89 *Analcellicampa* resequncing data using HAYSTAC^105^ with default parameters to detect the *Wolbachia* genome. Before examination, a custom database of complete *Wolbachia* genomes was compiled (Table S13), including 24 complete genomes from *Wolbachia* parasitizing Hymenoptera and other hosts, obtained from the NCBI RefSeq database, along with the *Wolbachia* genome assembled in this study. For samples with a higher number of confidently assigned *Wolbachia* reads (> 3000). We further ensured the results’ validity by using IGV^106^ to checking a detailed of the coverage and distribution of reads on *Wolbachia* genome. Both approaches provide robust support for confirming *Wolbachia* infection.

### Mapping and Variant Calling

We first performed quality control on raw sequencing reads using fastp^86^ for trimming and filtering. High-quality paired-end resequencing reads were then mapped to AD genome with BWA “BWA-MEM” algorithm^107^ using the default parameters. Post-alignment processing included sorting, marking duplicates, indel realignment, and base quality recalibration, which were carried out using Picard (http://picard.sourceforge.net, last accessed January 5, 2021) and Genome Analysis Toolkit (GATK)^108^ software. Genomic variants for each individual were then identified using the HaplotypeCaller module and the GVCF model in GATK. followed by merging all GVCF files. SNPs were called with the HaplotypeCaller module. We then kept SNP sites if they passed the recommended hard filtering thresholds (QD > 2, FS < 60, MQ > 40, MQRankSum > −12.5, and ReadPosRankSum > 15) as used previously^109^. Additional filtering was applied, retaining SNPs with a minor allele frequency (MAF) > 0.01 and SNP call rates > 80%. The resulting high-quality variants were used for downstream analyses. The transition-to-transversion (Ts/Tv) ratio was calculated via VCFtools^110^ to validate the quality of our SNPs set. Called genotypes were phased using SHAPEIT4^111^. SnpEff (v5.0)^112^ was used for annotating and predicting the effects of identified SNPs.

### Phylogenetic and Population Genetic Analyses

A maximum-likelihood tree was constructed using iqtree program (version: 1.6.12)^113^ in ASC model sites with optimal substitution model determined by the software and with 1,000 bootstrap replicates. Principal Component Analysis (PCA) was conducted using the “SNPRelate” package in R^114^ based on whole-genome SNPs dataset. ADMIXTURE [v1.3.0]^115^ was used to quantify the genome wide admixtures among all samples. Admixture was run for each possible genetic clusters (K = 2 to 9) with 10 replicates for each K. We ran CLUMPAK^116^ on ADMIXTURE outputs to generate consensus solutions for each K and finally identified the optimal K cluster supported by the data based on cross validation (CV) errors. Population splitting and admixture analyses were further carried out using TreeMix program^117^. This method requires unlinked markers, so we used SNPrelate to prune out markers with high linkage (r^2^ ≥ 0.2 within 100 Kb). We selected an appropriate number of migration edges based on the decay in variation explanation improvements with successive numbers. BITE^118^ was used to do 100 bootstrap replicates, and to visualize the consensus trees with bootstrap values and migration edges when doing TreeMix analyses. Identical scores (ISs) were calculated to evaluate the similarities of the sequenced genomes to the AD reference genome according to the following formula:

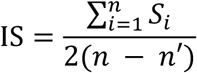

where *S_i_* is the number of alleles identical to the AD reference allele at a given SNP site *i*, *n* is the total number of SNPs within a 50-kb window and *n′* is the total number of missing SNPs within a 50-kb windows.

### Demographic History and Selection Analyses

We used PSMC^38^ to estimate variation in *Ne* over historical time. Diploid consensus sequences were generated using the ’mpileup’ command from the SAMtools (http://samtools.sourceforge.net/) package, applying the recommended PSMC settings (-C 50, -O, -D 2*reads_depth, -d 1/3*reads_depth). Input files were created using the ’fq2psmcfa’ tool. For each sample, the PSMC algorithm was run for 30 iterations. We estimated piecewise constant ancestral *Ne* over 90 time intervals, following the methodology of Wallberg et al^119^. To account for variance in *Ne*, we performed 100 bootstrap replications for representative samples from each subspecies. These steps ensured robust validation of historical population size fluctuations.

Since PSMC analyzes a single individual per population, we also employed SMC++ (v1.15.2)^41^ to infer population size dynamics using multiple individuals per population. To ensure robust mapping, uniquely mappable regions were identified using the SNPable toolkit (http://lh3lh3.users.sourceforge.net/snpable.shtml) with settings -k=35 and r=0.9. When analyzing population size dynamics separately for males and females, we focused on populations with more than one individual for each sex (AD, AX, and AM). As all individuals had a sequencing coverage exceeding 20-fold, there was no need for sample exclusion due to low coverage. Input files for SMC++ were prepared following the recommended pipeline from the SMC++ GitHub repository (https://github.com/popgenmethods/smcpp). All analyses were conducted using default parameters to ensure consistency across datasets.

Finally, we used momi2^37^, which fits models to the site-frequency spectrum, to further assess the underlying population history based on six populations (AD, AW, AX, AM, AA, and AE) and Outgroup, assuming a mutation rate of 5.27 × 10^-9^ per site per generation^119^ and a generation time of 1 year. Given that our study primarily focuses on changes in *Ne* across species, we tested two models: one assuming constant population size and another allowing for variable population size over the past 30,000 years, reflecting potential effects of the LGM and corresponding to the *Ne* declines observed during this period in PSMC analyses (Figure 2H and S8). The best-fitting model was selected based on the Akaike Information Criterion (AIC) value. Then, we performed 20 independent runs with different starting parameters and kept the model with the biggest log-likelihood value for the optimal model. To ensure robust inference of population parameters, we conducted 50 bootstraps, providing confidence intervals for the estimated values.

Nucleotide diversity (π) and dxy values were calculated for six species in and between *Analcellicampa* genus using pixy [v1.2.7]^120^ with a 10-kb window and a 5-kb sliding window. The average π or dxy was calculated using the formula: sum(window count_diffs)/sum(window comparisons). To detect genomic regions under selection in AD, we calculated the log_2_(π-_AD_/π-_AX_). We alsp used updated XP-CLR^121^ to compare the allele frequency distributions for the AD and AX populations to detect selective sweeps based on phased genotypes that were inferred by SHAPEIT4. The command line was *xpclr --input input.vcf --samplesA AD_list --samplesB AX_list –chr chrN –phased --size 10000 --step 5000 -maxsnps 600 –out outputFile*. The parameters used included overlapping sliding windows of 10 kb with a 5 kb step and a maximum of 600 SNPs per window. Genomic regions falling within the top 5% of XP-CLR values and log_2_(π ratio) values across the whole genome were identified as candidate selective sweeps. We carried out Gene Ontology (GO) enrichment analysis on the PSGs within candidate selective sweep regions using the enrichGO function from clusterProfiler [v3.14.3]^122^ and the org.Dm.eg.db database.

## Author Contributions

M.W. designed research; G.N., B.T. and D.W. contributed samples; M.Z. performed research; M.Z., R.Z., G.M. and J.Q. analyzed data; M.Z. and M.W. wrote the paper.

## Competing Interest Statement

The authors declare no competing interest.

## Acknowledgments

This work was financially by grants from National Natural Science Foundation of China (Grant No. 32370500), Jiangxi Provincial Natural Science Foundation (Grant No. 20232BAB215017), and Science and Technology Research Project of Jiangxi Provincial Department of Education (Grant No. GJJ2200344).

## Notes

### Competing Interest Statement

The authors have declared no competing interest.

