## Supplemental Figures for "Telomere-to-Telomere Genome Assembly Uncovers *Wolbachia*-Driven Sex-Specific Demography and Challenges Fisher’s Principle in a Sawfly"

Mingpeng Zhang et al.

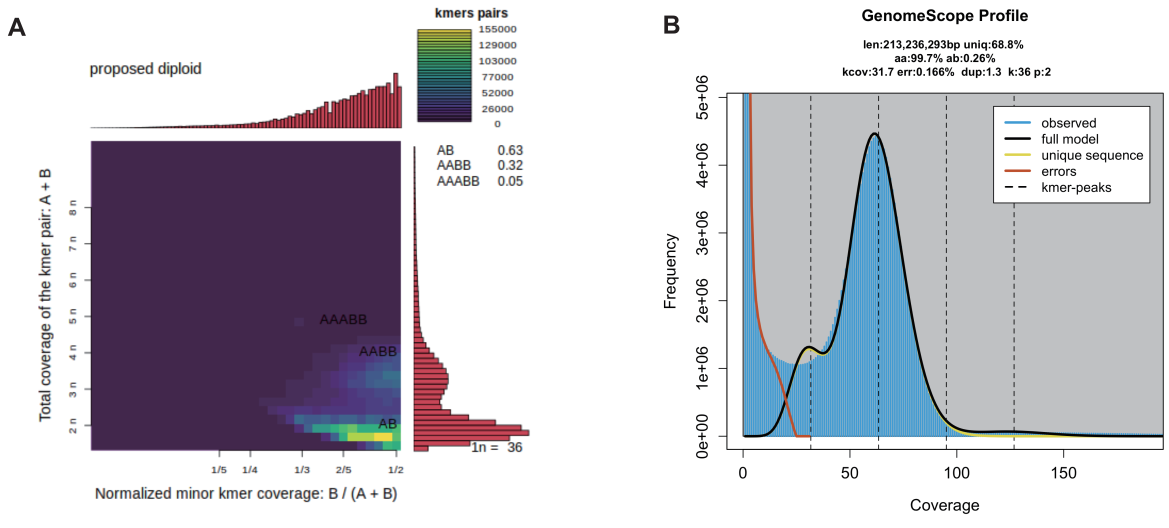


Figure S1. Genome profiling plots (A) shows ploidy assessment using Smudgeplot, and (B) shows estimated genome size and complexity as output by GenomeScope2.

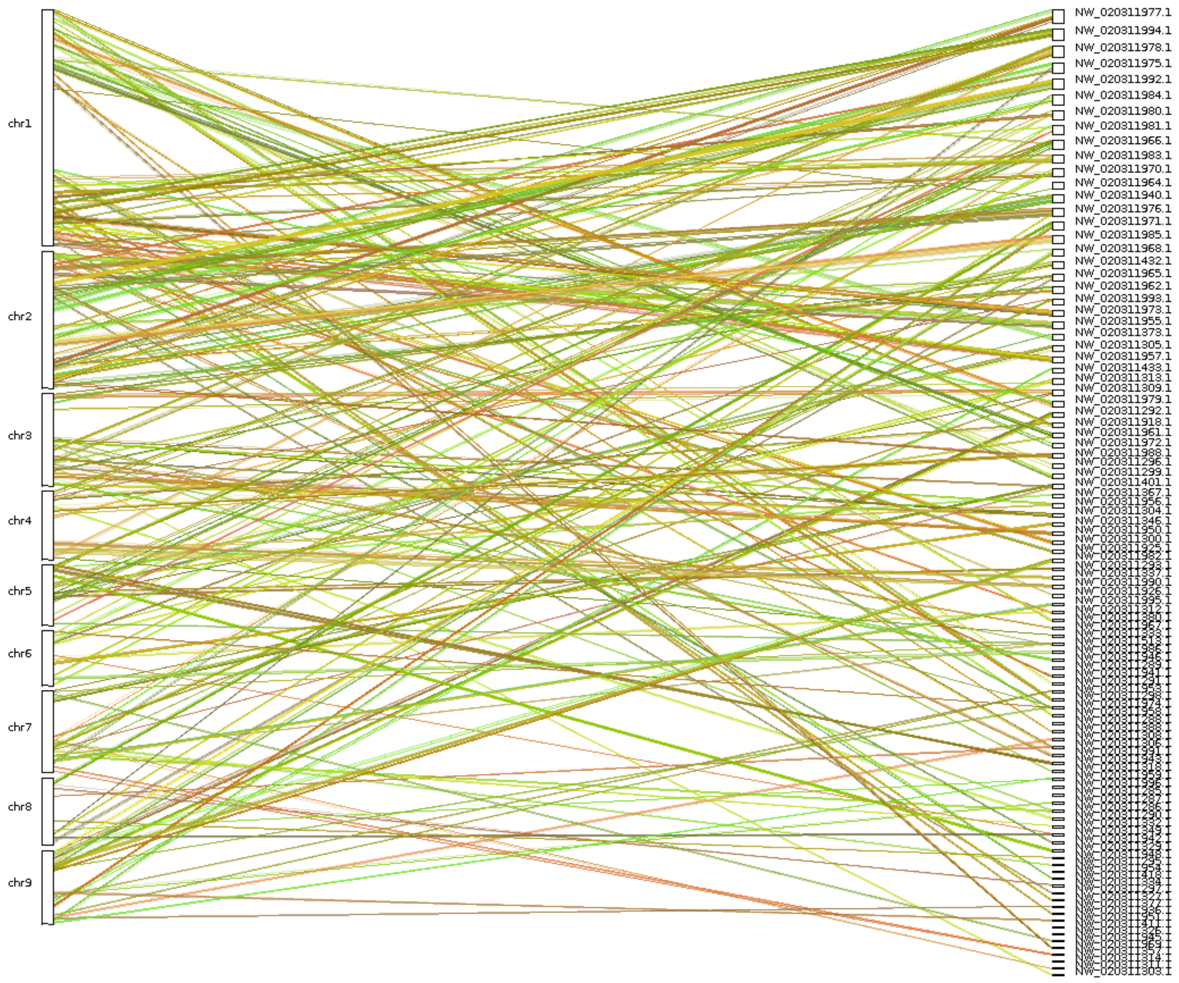


Figure S2. Dual synteny plot generated using MCScanX software. Chromosomes/scaffolds are labeled accordingly: *A.danfengensis* (left) and *Orussidae abietinus* (right).

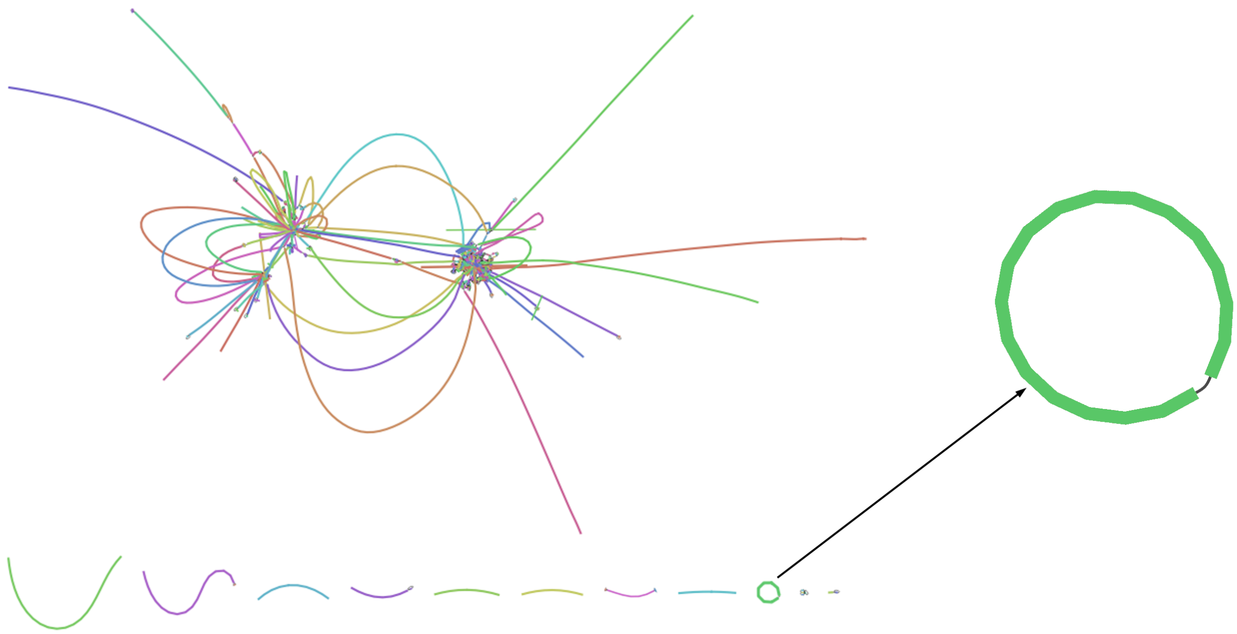

Figure S3. Assembly graph of the initial assembly generated by Flye, visualized with Bandage. The circular structure represents the *Wolbachia* genome.

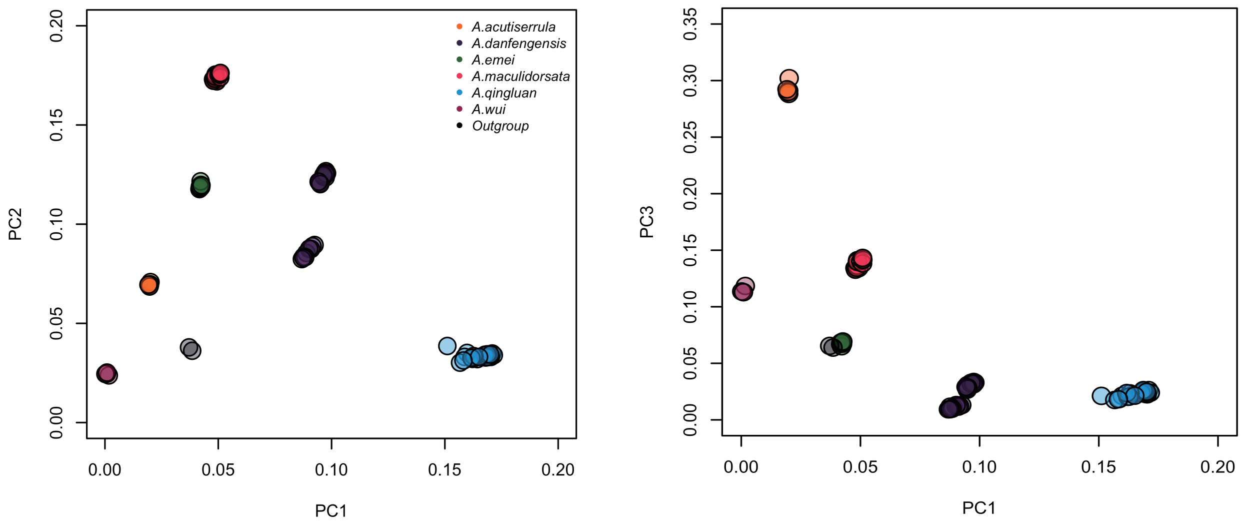


Figure S4. Principal component (PC) analysis plots based on the first three PCs.

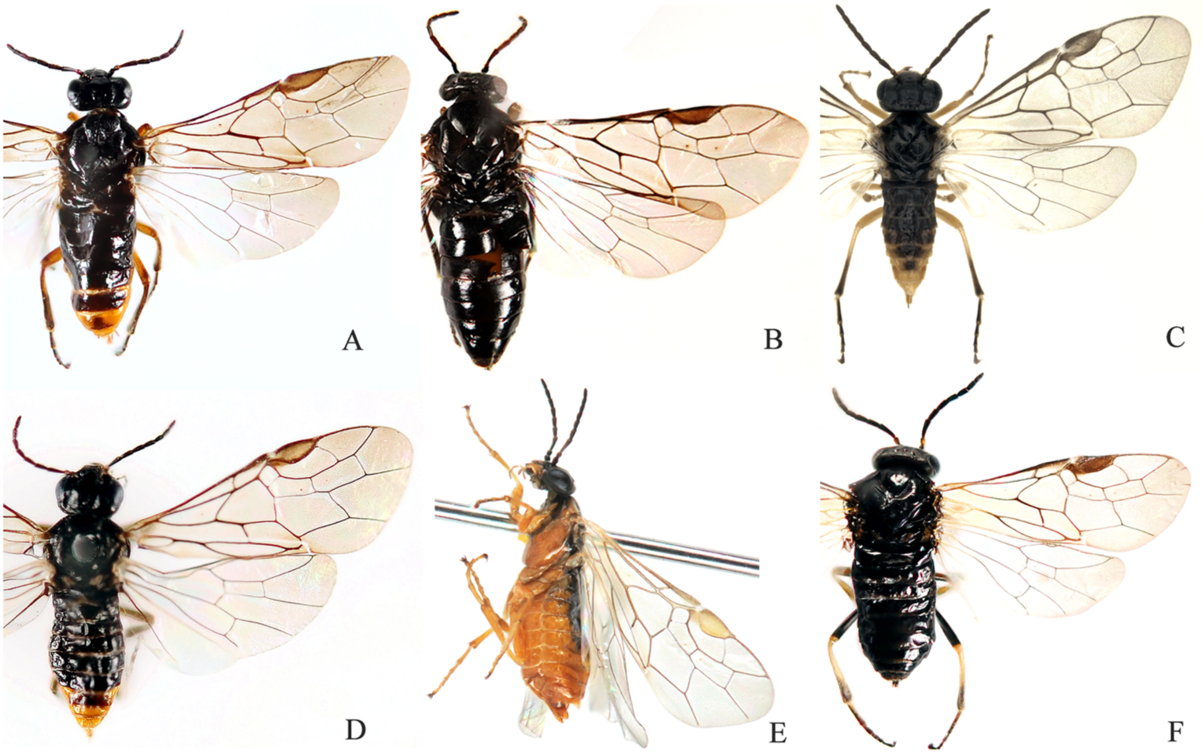


Figure S5. Pictures of *Analcellicampa* spp. Female.
A. *A. acutiserrula*; B. *A. danfengensis*; C. *A. emei*; D. *A. maculidorsata*; E. *A. xanthosoma*; F. *A. wui*

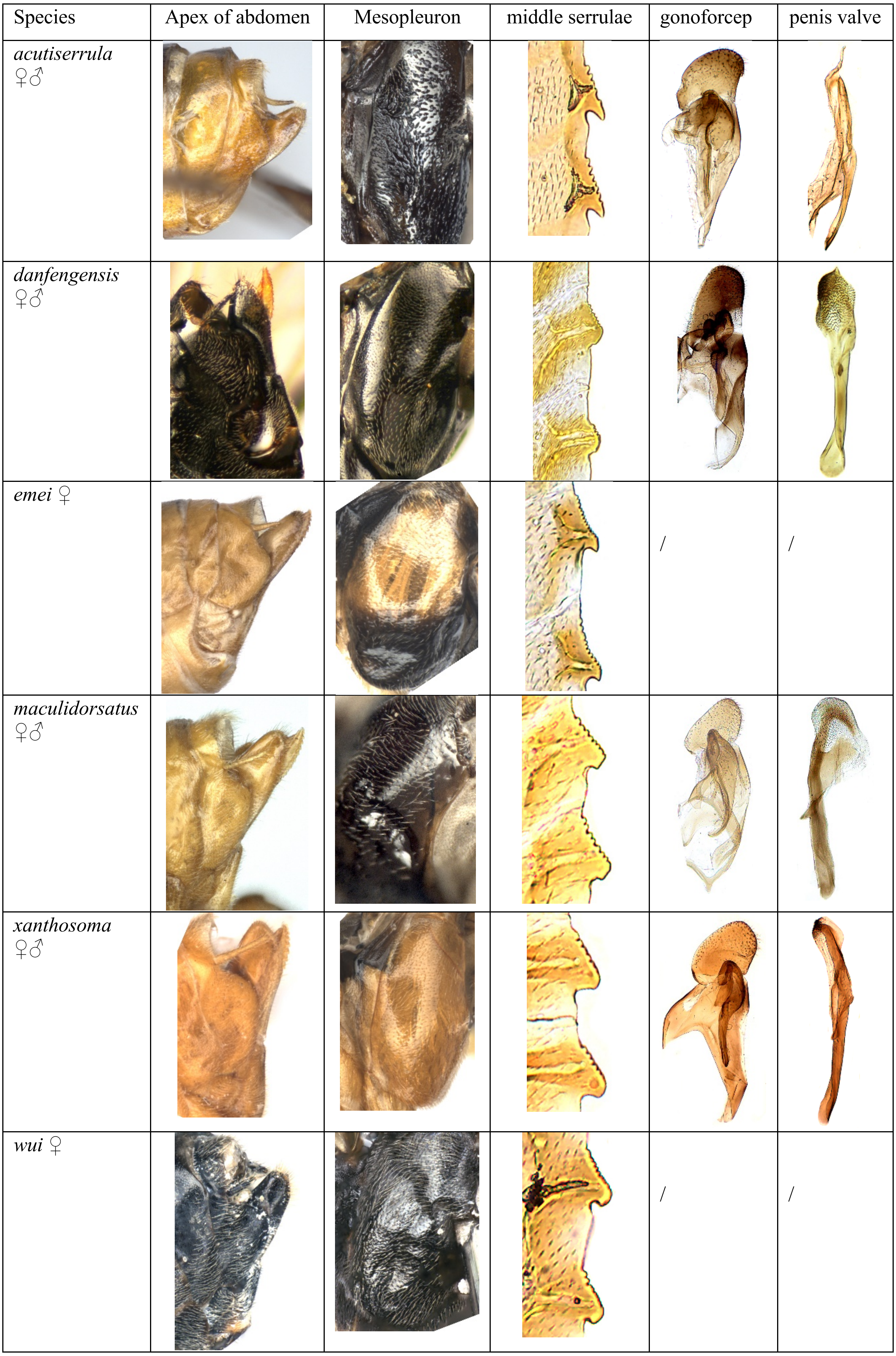


Figure S6. Partial morphological differences of *Analcellicampa* spp.

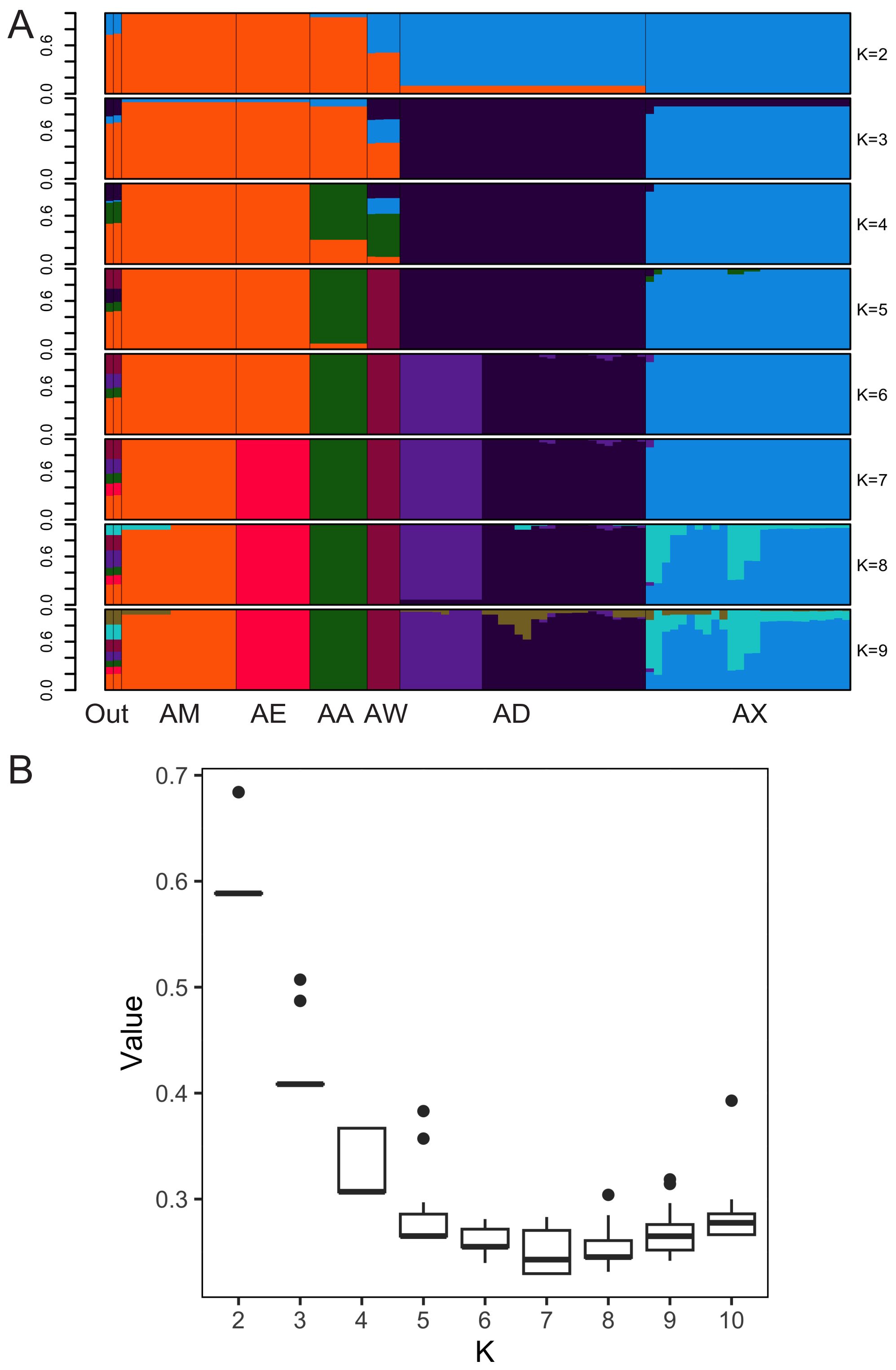


Figure S7. (A) ADMIXTURE analysis with K = 2–9. Colors in each column represent ancestry proportion. (B) The optimal K value with the lowest cross-validation (CV) error was 7 in ADMIXTURE analyses.

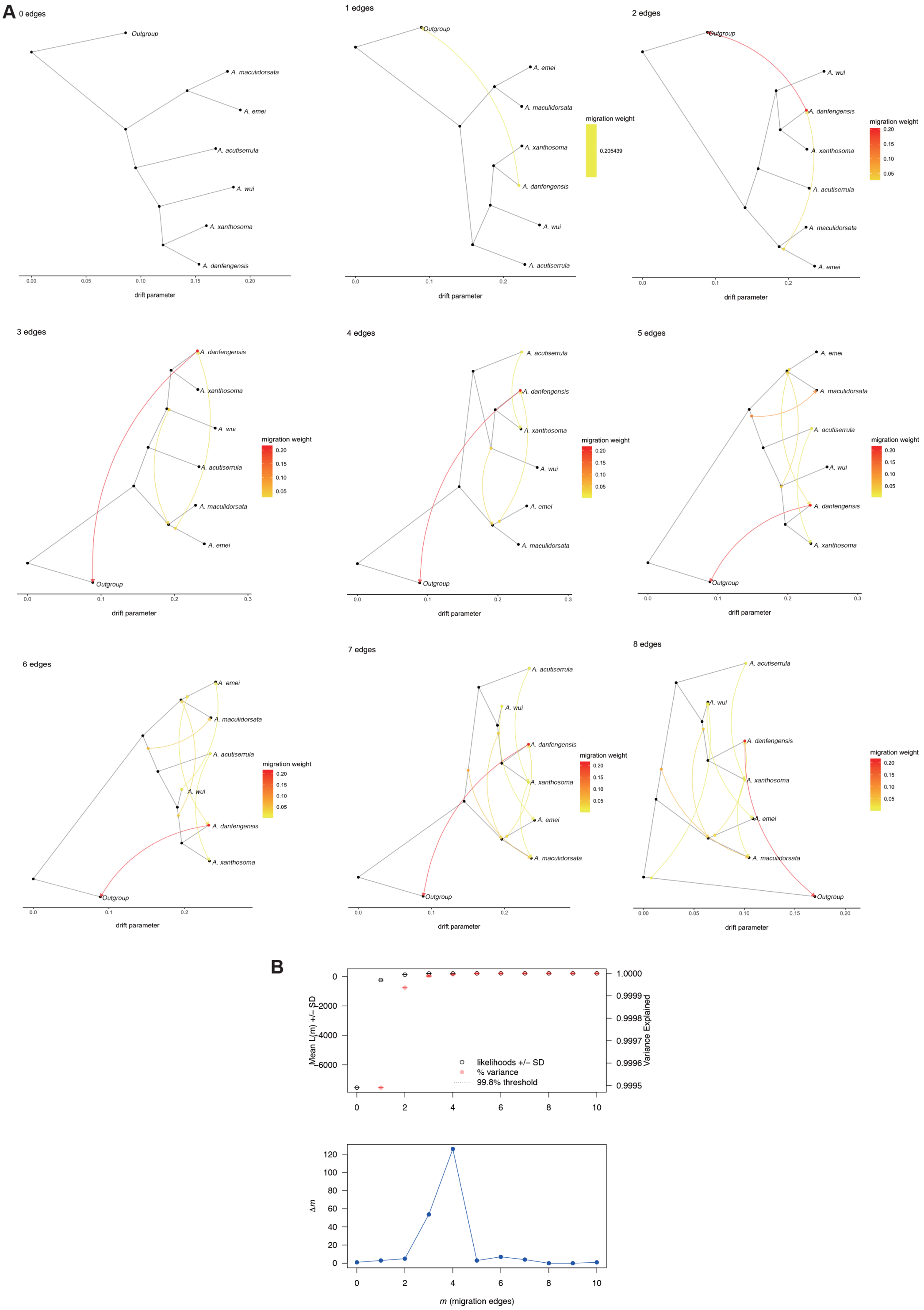


Figure S8. TreeMix results summary. (A) Phylogenetic trees and migration edges inferred with TreeMix for 0 to 8 migration edges. Migration edges are represented as arrows, indicating the direction of inferred gene flow between populations. (B) Changes in variation explained as additional migration edges are incorporated, illustrating the diminishing returns in explanatory power beyond four migration edges.

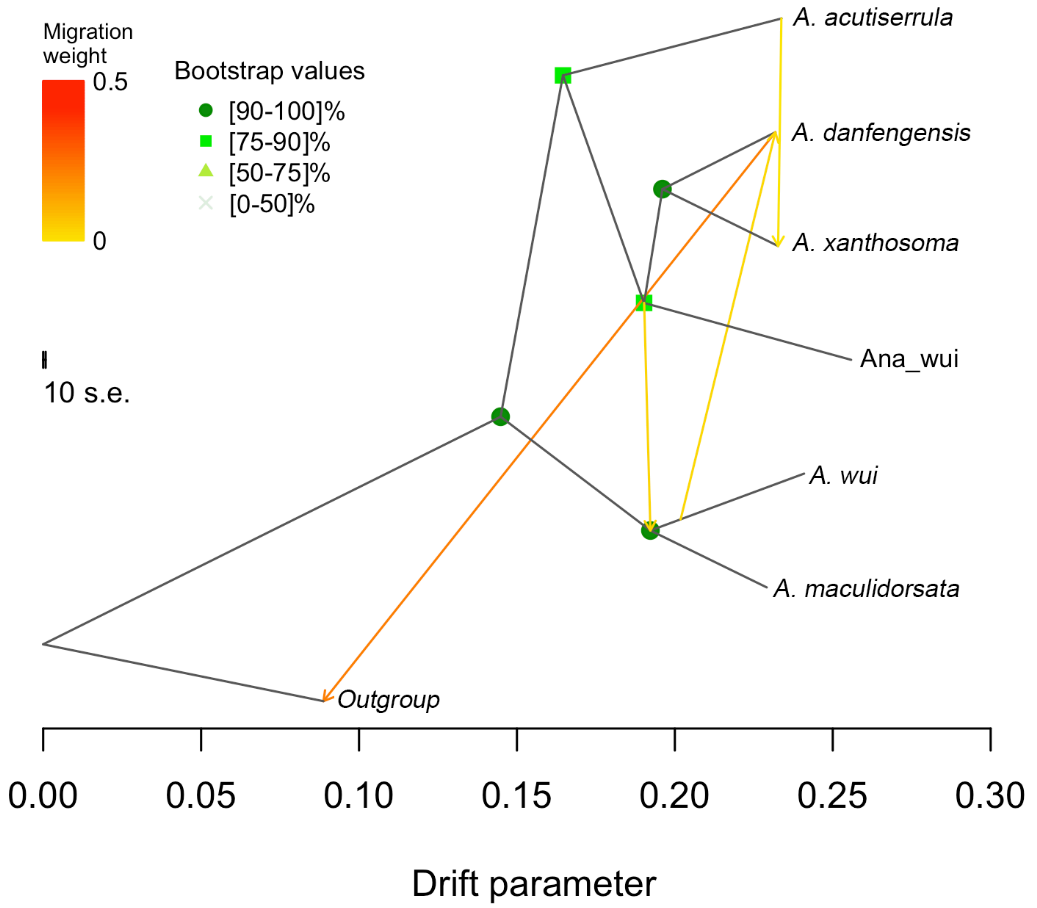


Figure S9. Optimal results inferred using TreeMix with four migration edges. Arrows indicate directions of inferred gene flow between populations, and branch lengths reflect genetic drift. Bootstrap values are shown for key nodes to indicate statistical support.

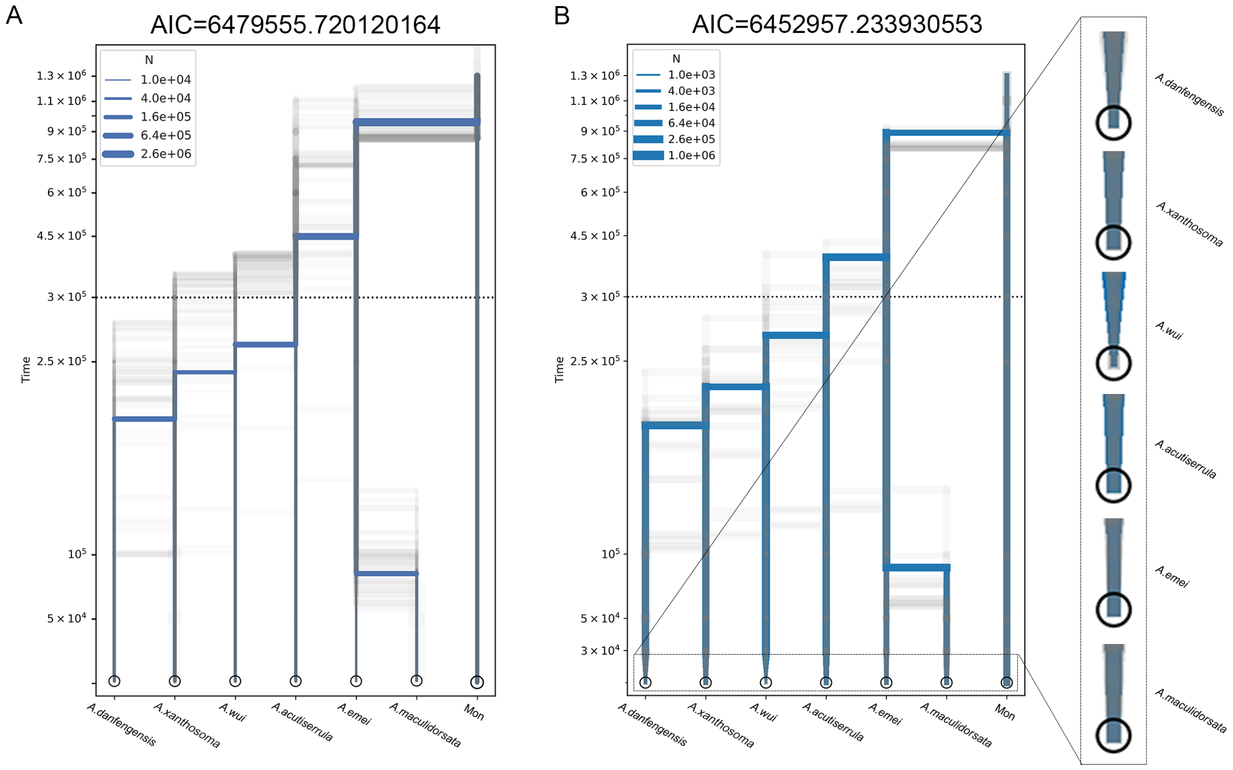


Figure S10. Momi2 Demographic Models: Model 1: Assumes constant population size over time. Model 2: Allows for variable population size dynamics from the Last Glacial Maximum (LGM) to the present.

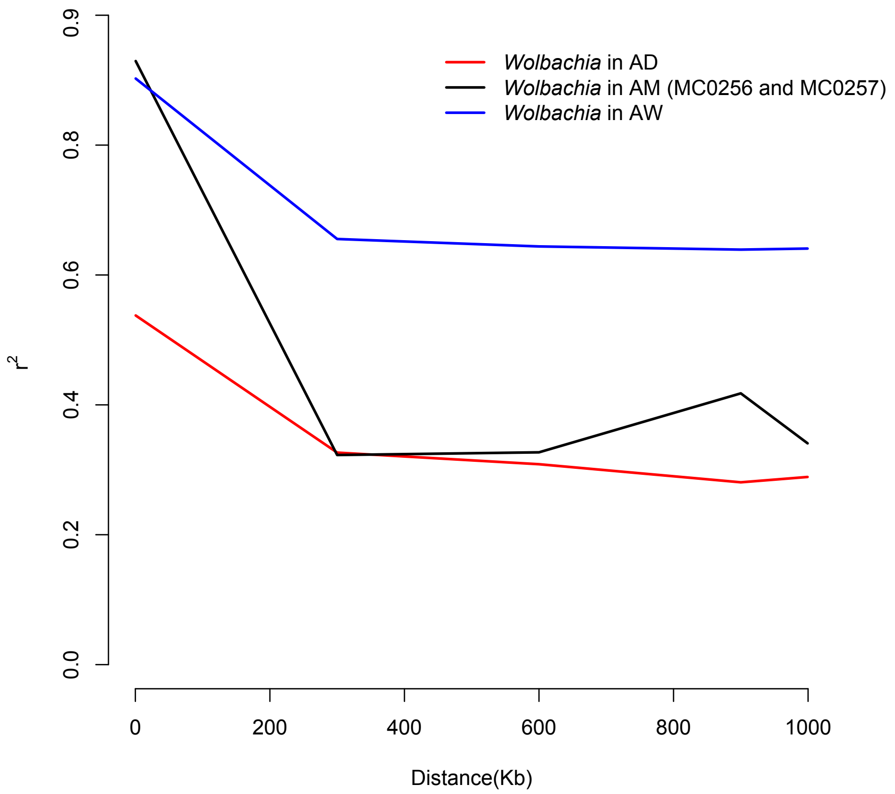


Figure S11. LD decay of three types of Wolbachia strains.

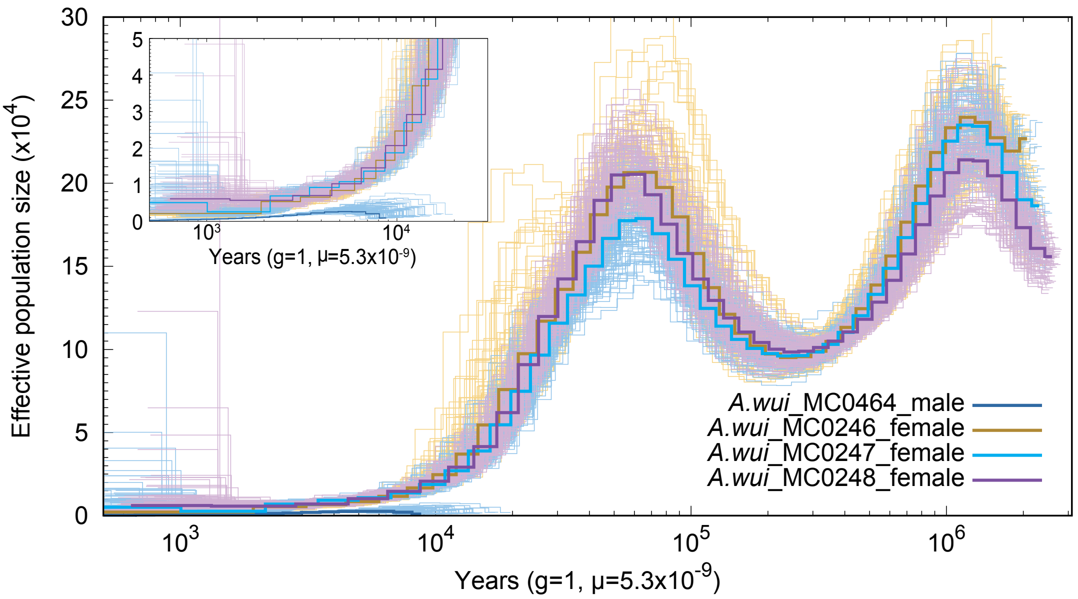


Figure S12. Demographic history inferred by PSMC shows the changes in effective population size of *A.wui* over past years.
